# Chalcogen derivatives for the treatment of African trypanosomiasis: biological evaluation of thio and seleno- semicarbazones and their azole derivatives

**DOI:** 10.1101/2025.02.21.639579

**Authors:** Mercedes Rubio-Hernández, Thaiz R. Teixeira, Tina P. Nguyen, Mai Shingyoji, Elany Barbosa da Silva, Anthony J. O’Donoghue, Conor R. Caffrey, Silvia Pérez-Silanes, Nuria Martínez

**Author notes:** (SPS); (NM); (CRC). SPS, NM and CRC contributed equally to this work.

## Abstract

Human African Trypanosomiasis (HAT) is caused by *Trypanosoma brucei*. Drug therapy remains challenging due to drug resistance and/or toxicity. New drugs are needed. Using thiosemicarbazones as a starting point, we employed a *S* to *Se* isosteric replacement strategy to design 44 analogs which were evaluated against *T. brucei in vitro*. Compounds were divided into eleven groups of four derivatives corresponding to thio-, selenosemicarbazones, and their cyclic counterparts, thio- and selenazoles. We selected three groups which contained a total of six derivatives that inhibited parasite growth by >70%. Then, we investigated the mechanism of action of these compounds, performing quantitative assays to measure their inhibition of the *T. brucei* cathepsin L-like protease (*Tbr*CATL) and DPPH antioxidant activities. The lead compound (***Se*O3**) showed antioxidant capacity and the best activity against *T. brucei* (EC_50_ = 0.47 µM). Nevertheless, its toxicity should be improved. We also predicted the interactions of these compounds with *Tbr*CATL utilizing molecular dynamics. We demonstrate that the *Se* derivatives are more active than their *S* analogues, and that the selenazole ring decreases *Se*-associated toxicity. Also, thio- and selenosemicarbazones are more potent against *Tbr*CATL than the cyclic derivatives. We conclude that *Tbr*CATL inhibition should be combined with antioxidant activity to obtain active compounds against *T. brucei*.

## INTRODUCTION

Human African trypanosomiasis (HAT; aka sleeping sickness) is a neglected tropical disease (NTD) that is caused by *Trypanosoma brucei*, a protozoan parasite that infects humans in Sub-Saharan Africa. HAT is caused by two subspecies of *T. brucei*, *Trypanosoma brucei gambiense* and *Trypanosoma brucei rhodesiense*, which are transmitted by the bite of the tsetse fly (*Glossina* spp.). Most cases (92%) are due to *T. b. gambiense*, which is prevalent in west and central of Africa, and develops as a chronic, and ultimately, fatal disease that affects the central nervous system (CNS). *T. b. rhodesiense* (8% of cases) is endemic in southern and eastern parts of Africa, and causes a more acute, but likewise, fatal, CNS disease [1,2]. In livestock, particularly cattle, *Trypanosoma brucei brucei* is one of a number of African trypanosomes responsible for *Nagana* disease [3], while also being used as a laboratory model for both of the human-infective *T. brucei* sub-species.

To treat HAT, the few medicines available suffer from toxicity and administration routes that complicate treatment (intramuscular for pentamidine or intravenous for melarsoprol, suramin and eflornithine) in resource-poor medical environments [1]. Moreover, the choice of drug depends on the stage of the disease diagnosed and the causative agent. Specifically, pentamidine and suramin are used for first (hemolymphatic) stage infections of *T. b. gambiense* and *T. b. rhodesiense*, respectively; whereas melarsoprol, and nifurtimox combined with eflornithine, are utilized to treat the second (CNS-infiltrated) stage of disease due to *T. b. gambiense* and *T. b. rhodesiense*, respectively [4]. Resistance, including cross-resistance, has been documented for melarsoprol [5–7], pentamidine [5] and eflornithine [6]. Since its approval in 2018, the oral drug, fexinidazole, has been recommended for both stages of the disease regardless of infecting subspecies, however, this drug has side effects [8,9]. Overall, there is a need to discover new orally-administered alternatives with acceptable safety profiles.

Recent advances in the search for pharmacological therapies have included the design of inhibitors of a cysteine protease known as *T. brucei* cathepsin L (*Tbr*CATL; formerly known as rhodesain [10,11]). *Tbr*CATL is the major papain family, cysteine protease found in *T. brucei* [12]. This enzyme contributes to the parasite’s ability to cross the blood-brain barrier (BBB) that facilitates the second stage of HAT [13]. The protease also participates in various processes [14], such as nutrition [15], host cell invasion [16] and differentiation [17]. *Tbr*CATL is homologous to cruzain (Cz) in *Trypanosoma cruzi*, the etiological agent of Chagas disease [18]. Cz and *Tbr*CATL are validated drug targets [19,20] and share 70% of the sequence, and differing by just two amino acids in the active site [18,21]. Therefore, compounds that inhibit Cz and *T. cruzi* growth [22,23], might also demonstrate activity against *Tbr*CATL and *T. brucei*. Previously, synthetic compounds inhibiting *Tbr*CATL by >99% have shown trypanocidal activity [12,14].

Thiosemicarbazones (*S*-semicarbazones) and thiazoles have demonstrated activity against *T. brucei* and inhibit *Tbr*CATL [21,24–27]. *S*-semicarbazones are particularly suitable for the treatment of NTDs because of their low-cost synthesis, low molecular weight and nonpeptidic nature. They are also covalent reversible inhibitors of cysteine proteases, and serve as intermediates in the synthesis of thiazoles [26,28–31]. Given these benefits, we investigated the isosteric replacement strategy of changing sulfur (*S*) to selenium (*Se*) [32], to synthesize the *Se*-counterparts of *S*-semicarbazones and thiazoles, which are, respectively, selenosemicarbazones (*Se*-semicarbazones) and selenazoles. *S* and *Se* both belong to the chalcogen group, and they are bioisosteres, thus, their physicochemical properties are very similar. However, because *Se* is slightly bigger, the valence electrons are loosely bonded, increasing the reactivity of *Se* and making it more nucleophilic and polarizable compared to *S*. The increased reactivity of *Se* contributes to its electronegativity and redox properties [33,34], which are imparted to the molecules that contain this element [35–37].

*Se* is a trace element essential for both parasite and human survival, however, overdosing can lead to toxicity. This element stimulates the immune system and is a key component in selenoproteins, such as glutathione peroxidase and selenocysteine [34]. *Se* deficiency does not directly cause disease but weakens the organism to facilitate various pathological conditions [33,38,39]. Compounds containing *Se* have been explored to treat different parasitoses like Chagas disease [38,40] and leishmaniasis [41]. However, data for *Se*-containing small molecules against *T. brucei* are scarce. To date, only the anti-inflammatory and anti-oxidant compound, ebselen, is known to inhibit *T. brucei* hexokinase 1 [42,43] and trypanothione synthetase [44], two enzymes that participate in glycolysis [42] and in the defense against the oxidative damage [44], respectively. In addition, an *in vivo* study showed that *T. brucei*-infected rats improved their response to the disease when supplemented with dietary *Se* [33]. Consequently, we consider the isosteric replacement of *S* with *Se* an interesting strategy in the pursuit of new therapies to treat HAT.

In this work, we evaluated two series of compounds that were previously shown to kill *T. cruzi* (0.31 – 3.73 µM) and inhibit Cz (1.56 – 81.69 nM) [40]. The first series contained “open” *S*-semicarbazone (***S*O**) and *Se*-semicarbazone (***Se*O**) derivatives. The second series comprised the corresponding cyclic or “closed”, thiazole (***S*C**) and selenazole (***Se*C**) counterparts (**Figure 1**). With these series, our objective here was to identify a compound(s) that selectively kill *T. brucei* and inhibit *Tbr*CATL.

**Figure 1.**
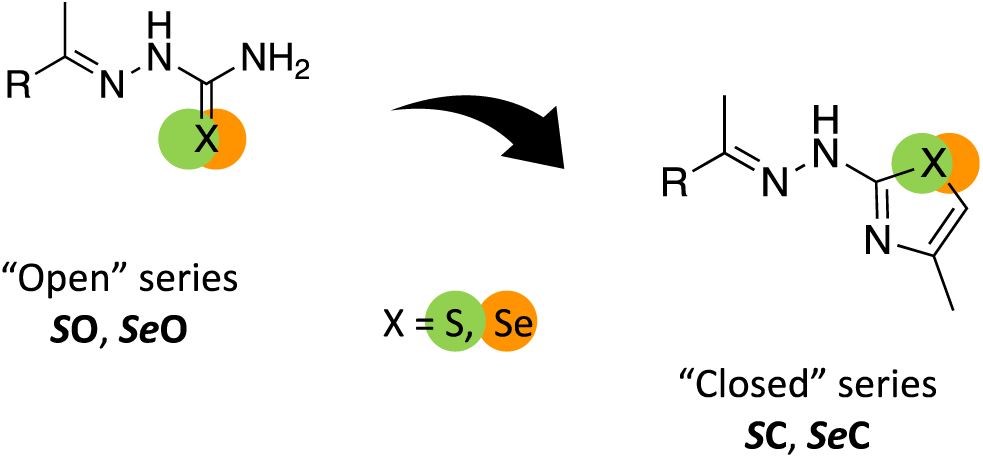
General structures of the compounds studied: *S*-semicarbazones (***S*O**), *Se*-semicarbazones (***Se*O**), thiazoles (***S*C**) and selenazoles (***Se*C**).

## RESULTS

### Biological evaluation

The activity of 44 compounds (***S*O1*-S*O11**, ***Se*O1-*Se*O11**, ***S*C1-*S*C11**, ***Se*C1-*Se*C11**) was evaluated against *T. brucei brucei* Lister 427 *in vitro* [45]. First pass screens at 10 µM identified 12 actives (Supporting **Table S1**) that inhibited parasite growth by >70%. These 12 compounds were selected for dose response (DR) analysis, as well as for DR counter cytotoxicity assays using human embryonic kidney (HEK)293 and hepatoblastoma (Hep)G2 cells (Supporting **Table S1**), both of which are commonly used to assess compound toxicity. All 12 compounds showed >50% cytotoxicity for both cell lines (Supporting **Table S1**), but the concentrations at which cell growth was inhibited by 50% (CC_50_ values) were at least twice as high for the selenazoles as for the *Se*-semicarbazones (Supporting **Table 1**). ***Se*O3** was the most potent compound with an EC_50_ value (concentration at which parasite growth is inhibited by 50%) of 0.47 µM. Relative to the CC_50_ values of 2.70 and 2.82 µM obtained with the HEK293 and HepG2 cells, respectively, the selectivity index (SI) for ***Se*O3** was the highest of the compounds tested with a value of approximately 6.

Based on the parasiticidal and toxicity data, we selected the *Se*-compounds, ***Se*O1**, ***Se*C1**, ***Se*O3**, ***Se*C3**, ***Se*O5**, ***Se*C5** for additional assays (see below). We also included the corresponding *S*-counterparts (***S*O1**, ***S*C1**, ***S*O3**, ***S*C3**, ***S*O5**, ***S*C5**) for comparison. **Table 1** shows the biological data for this top 12 hits.

**Table 1.**
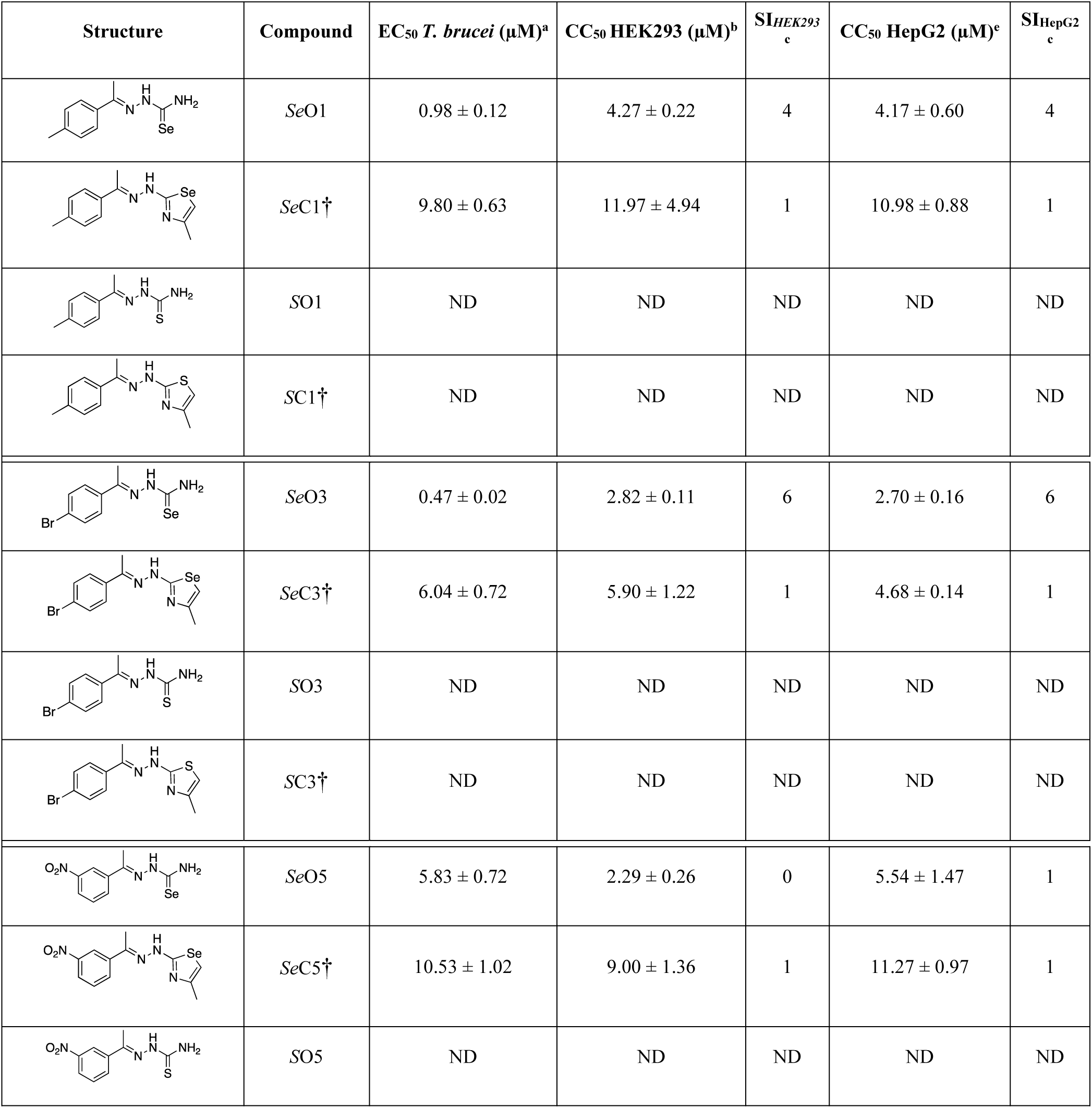
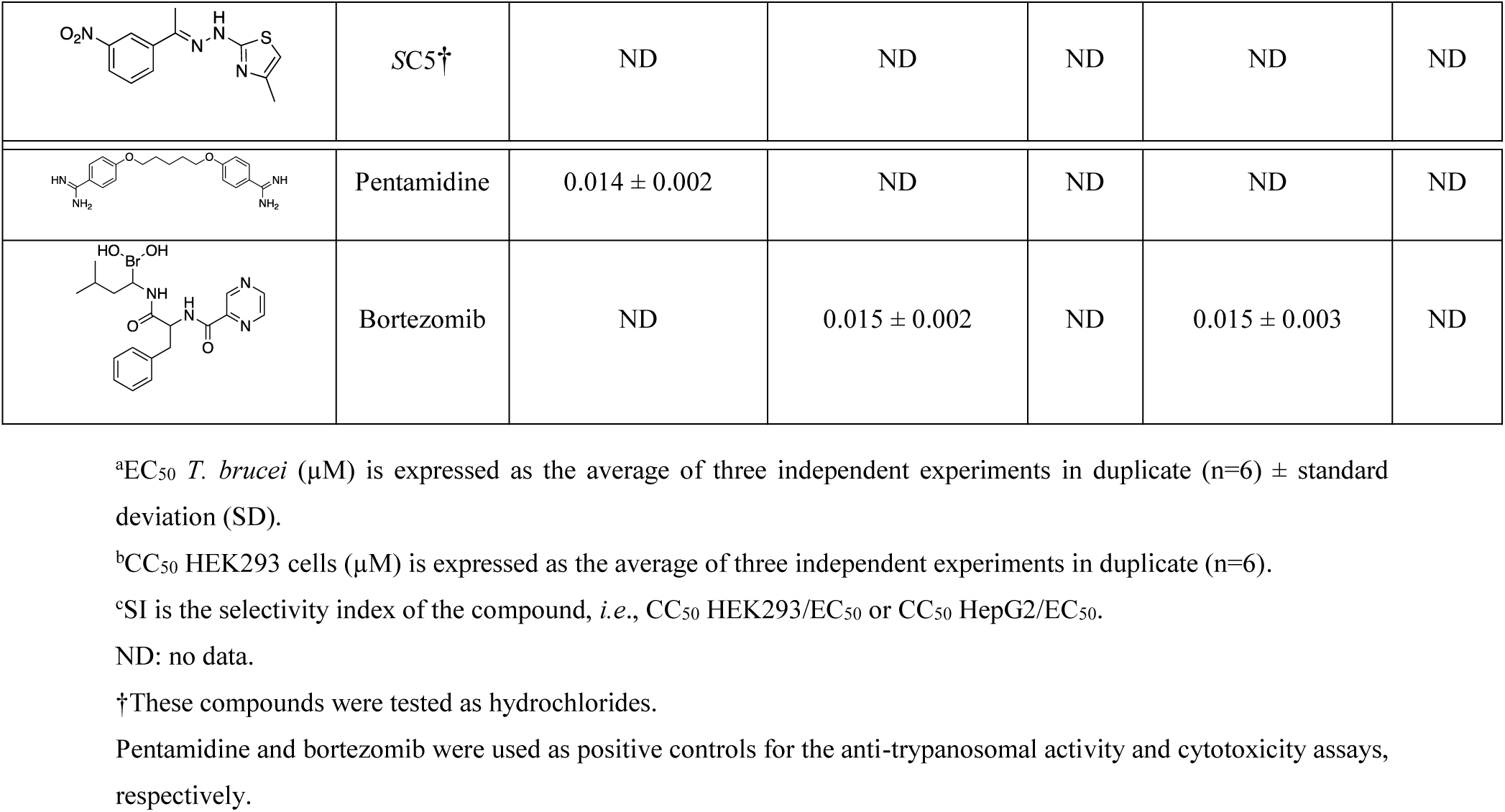
Biological data for the top 12 hits against *T. brucei*, and HEK293 and HepG2 cells.

| Structure | Compound | EC<sub>50</sub> *T. brucei* (μM)<sup>a</sup> | CC<sub>50</sub> HEK293 (μM)<sup>b</sup> | SI<sub>HEK293</sub><sup>c</sup> | CC<sub>50</sub> HepG2 (μM)<sup>e</sup> | SI<sub>HepG2</sub><sup>c</sup> |
| --- | --- | --- | --- | --- | --- | --- |
|  | **SeO1** | 0.98 ± 0.12 | 4.27 ± 0.22 | 4 | 4.17 ± 0.60 | 4 |
|  | **SeC1**† | 9.80 ± 0.63 | 11.97 ± 4.94 | 1 | 10.98 ± 0.88 | 1 |
|  | **SO1** | ND | ND | ND | ND | ND |
|  | **SC1**† | ND | ND | ND | ND | ND |
|  | **SeO3** | 0.47 ± 0.02 | 2.82 ± 0.11 | 6 | 2.70 ± 0.16 | 6 |
|  | **SeC3**† | 6.04 ± 0.72 | 5.90 ± 1.22 | 1 | 4.68 ± 0.14 | 1 |
|  | **SO3** | ND | ND | ND | ND | ND |
|  | **SC3**† | ND | ND | ND | ND | ND |
|  | **SeO5** | 5.83 ± 0.72 | 2.29 ± 0.26 | 0 | 5.54 ± 1.47 | 1 |
|  | **SeC5**† | 10.53 ± 1.02 | 9.00 ± 1.36 | 1 | 11.27 ± 0.97 | 1 |
|  | **SO5** | ND | ND | ND | ND | ND |
|  | SC5<sup>†</sup> | ND | ND | ND | ND | ND |
|  | Pentamidine | 0.014 ± 0.002 | ND | ND | ND | ND |
|  | Bortezomib | ND | 0.015 ± 0.002 | ND | 0.015 ± 0.003 | ND |

<sup>a</sup>EC<sub>50</sub> *T. brucei* (μM) is expressed as the average of three independent experiments in duplicate (n=6) ± standard deviation (SD).
<sup>b</sup>CC<sub>50</sub> HEK293 cells (μM) is expressed as the average of three independent experiments in duplicate (n=6).
<sup>c</sup>SI is the selectivity index of the compound, *i.e.*, CC<sub>50</sub> HEK293/EC<sub>50</sub> or CC<sub>50</sub> HepG2/EC<sub>50</sub>.
ND: no data.
<sup>†</sup>These compounds were tested as hydrochlorides.
Pentamidine and bortezomib were used as positive controls for the anti-trypanosomal activity and cytotoxicity assays, respectively.

### Mechanism of action: Enzyme inhibition assays

We conducted enzyme inhibition assays with *Tbr*CATL and the selected top 12 compounds (**Table 1**) to gain deeper insight into their possible mechanism of action (See **Table 2**). These assays also enable a direct comparison of structural and functional differences between the *S* and *Se* compounds as well as the open and closed compounds. First pass screens at 10 µM identified eight compounds that inhibited the activity of *Tbr*CATL by >85% (**Table 2**). Of these eight, five were *Se-*compounds (***Se*O1**, ***Se*O3**, ***Se*C3**, ***Se*O5** and ***Se*C5**) and three were *S*-compounds (***S*O3**, ***S*O5** and ***S*C5**). These compounds were subjected to DR analysis to measure the IC_50_ values (*i.e*., the concentration of compound necessary to inhibit 50% of the enzyme’s activity). In parallel, and to assess compound selectivity for *Tbr*CATL, we measured their IC_50_ values for inhibition of the orthologous human cathepsin L (*h*CatL; **Table 2**). The DR data for the five *Se-*compounds against both enzymes are shown in **Figure 2**. We note the shift to the left of the IC_50_ values for the open compounds relative to their cyclic derivatives, especially for ***Se*O3** which was 100 and 300 times better than ***Se*C3** against *Tbr*CATL and *h*CatL, respectively. The DR data for the *S*-compounds shows the same shift to the left (**Figure S1)**. Nevertheless, the IC_50_ values of the compounds within the same series (***Se*O1**, ***Se*O3**, ***S*O3**, ***Se*O5**, ***S*O5**) are similar and independent of the chalcogen atom present in the molecule (**Table 2**). Based on the selectivity of inhibition (*i.e*., IC_50_ *h*CatL / IC_50_ *Tbr*CATL), the best compound was ***S*C5** (SI = 49), a *S-*derivative that does not have activity against *T. brucei*. Of the compounds shown in **Tables 1** and **2**, ***Se*O3** stood out for combining activity against *T. brucei* with inhibition of *Tbr*CATL (EC_50_ = 0.47 µM and SI*_Tbr_*_CATL_ *=* 8).

**Figure 2.**
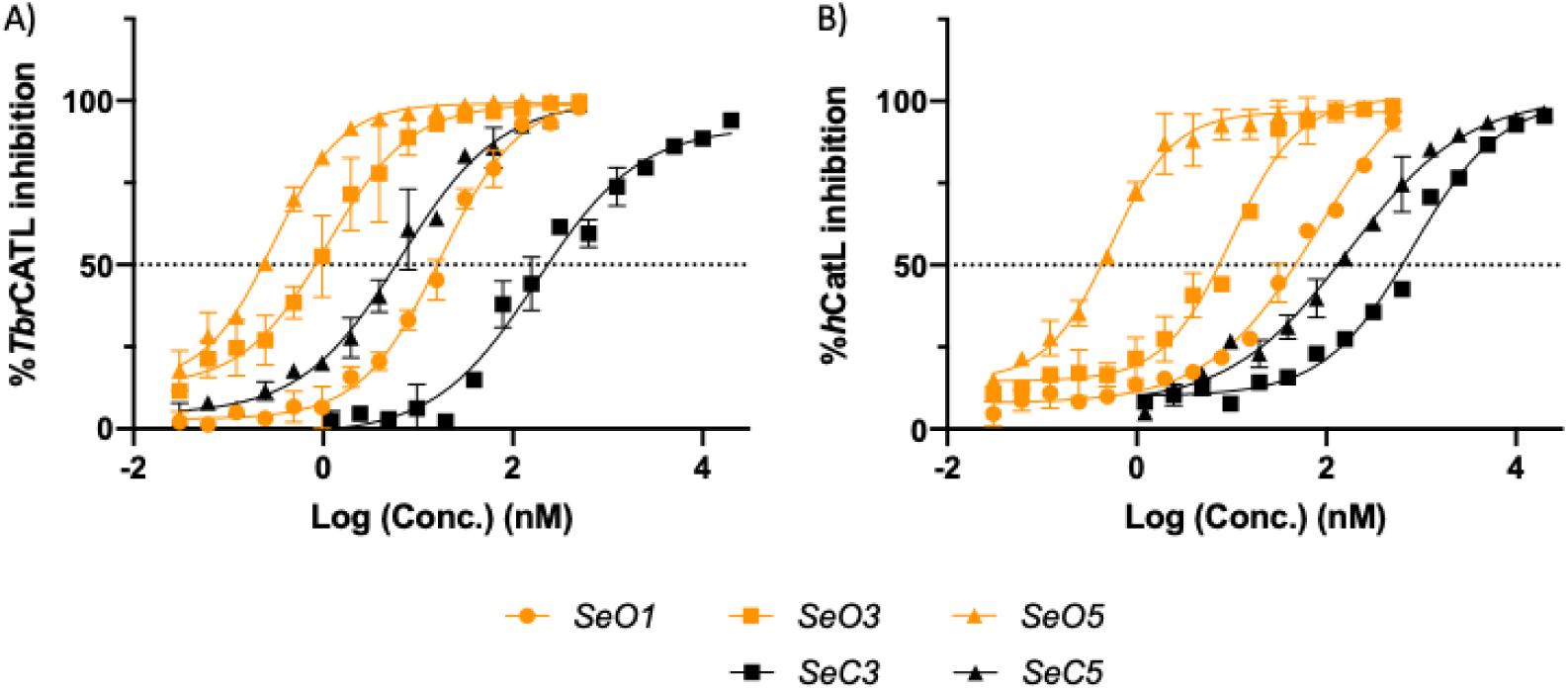
DR data for *Se* compounds against *Tbr*CATL (A) and *h*CatL (B).

**Table 2.** Inhibition of *Tbr*CATL and *h*CatL by the top 12 anti-trypanosomal agents.

| Compound | *Tbr*CATL inhibition (%)<sup>a</sup> | IC<sub>50</sub> *Tbr*CATL (nM)<sup>b</sup> | IC<sub>50</sub> *h*CatL (nM)<sup>b</sup> | SI<sub>*Tbr*CATL</sub><sup>c</sup> |
| --- | --- | --- | --- | --- |
| *Se*O1 | 97.08 | 17.90 ± 3.62 | 84.52 ± 17.03 | 5 |
| *Se*C1† | 51.07 | ND | ND | ND |
| *SO*1 | 79.98 | ND | ND | ND |
| *SC*1† | 32.85 | ND | ND | ND |
| *Se*O3 | 99.15 | 1.22 ± 0.52 | 9.87 ± 0.63 | 8 |
| *Se*C3† | 101.02 | 201.65 ± 0.92 | 792.75 ± 66.54 | 4 |
| *SO*3 | 95.54 | 7.33 ± 0.27 | 135.95 ± 19.87 | 19 |
| *SC*3† | 53.99 | ND | ND | ND |
| *Se*O5 | 101.47 | 0.30 ± 0.07 | 0.51 ± 0.08 | 2 |
| *Se*C5† | 98.02 | 6.91 ± 2.64 | 174.70 ± 43.70 | 25 |
| *SO*5 | 100.00 | 5.83 ± 0.35 | 20.74 ± 4.36 | 4 |
| *SC*5† | 87.70 | 82.57 ± 4.05 | 4065.50 ± 382.54 | 49 |
| E-64 | ND | 3.1 ± 0.0* | 30.0 ± 4** | 10 |

<sup>a</sup>*Tbr*CATL inhibition at 10 μM is shown as a percentage (%).
<sup>b</sup>IC<sub>50</sub> (nM) is shown as the mean ± standard deviation (SD) from two different experiments, each performed in triplicate (n=6).
<sup>c</sup>SI<sub>*Tbr*CATL</sub> is the selectivity of inhibition of the compound (IC<sub>50</sub>) for *Tbr*CATL compared to the IC<sub>50</sub> value for *h*CatL.
ND: no data.
\*IC<sub>50</sub> value for inhibition of *Tbr*CATL by E-64 (from [46]).
\*\*IC<sub>50</sub> value for inhibition of *h*CatL by E-64 (from [47]).
†These compounds were tested in the hydrochloride form.

### Mechanism of action: Molecular dynamics (*Tbr*CATL)

To study the interactions of ***Se*O1**, ***Se*O3**, ***Se*O5**, ***Se*C3**, and ***Se*C5** with *Tbr*CATL, molecular dynamics (MD) simulations were conducted. Our aim in this assay is to better understand the differences in the interactions between the open and cyclic *Se-*compounds. The ligand, K11777 (a covalent irreversible cysteine protease inhibitor), in the 2P7U [48] crystal structure was replaced with the different *Se* derivatives prior to running the MD simulations. A 500 ns trajectory was produced and analyzed to evaluate the stability of the complexes. Root mean square deviation (RMSD) values and root mean square fluctuation (RMSF) profiles were obtained (**Figure 3**). RMSD analysis indicated that the protein reaches structural stabilization in the complex at different times during the simulations depending on the ligand. The *Tbr*CATL in complex with ***Se*O1** remained stable from 120 to 260 ns, with a mean RMSD value of 2.5 Å. In the brominated derivative ***Se*O3** complex, the protein required 150 ns to stabilize, exhibiting a RMSD value of 1.93 Å, whereas its closed analog, ***Se*C3**, the enzyme presented lower deviations from 60 ns to the end of the trajectory (1.19 Å). The protease in complex with ***Se*O5** was rapidly stabilized within 20 ns achieving a mean RMSD value of 1.09 Å, whereas with the closed nitro derivative, ***Se*C5**, stabilization occurred at 150 ns with a mean RMSD value of 1.94 Å. The RMSF profiles demonstrated a similar pattern of residue fluctuations across all five complexes.

**Figure 3.**
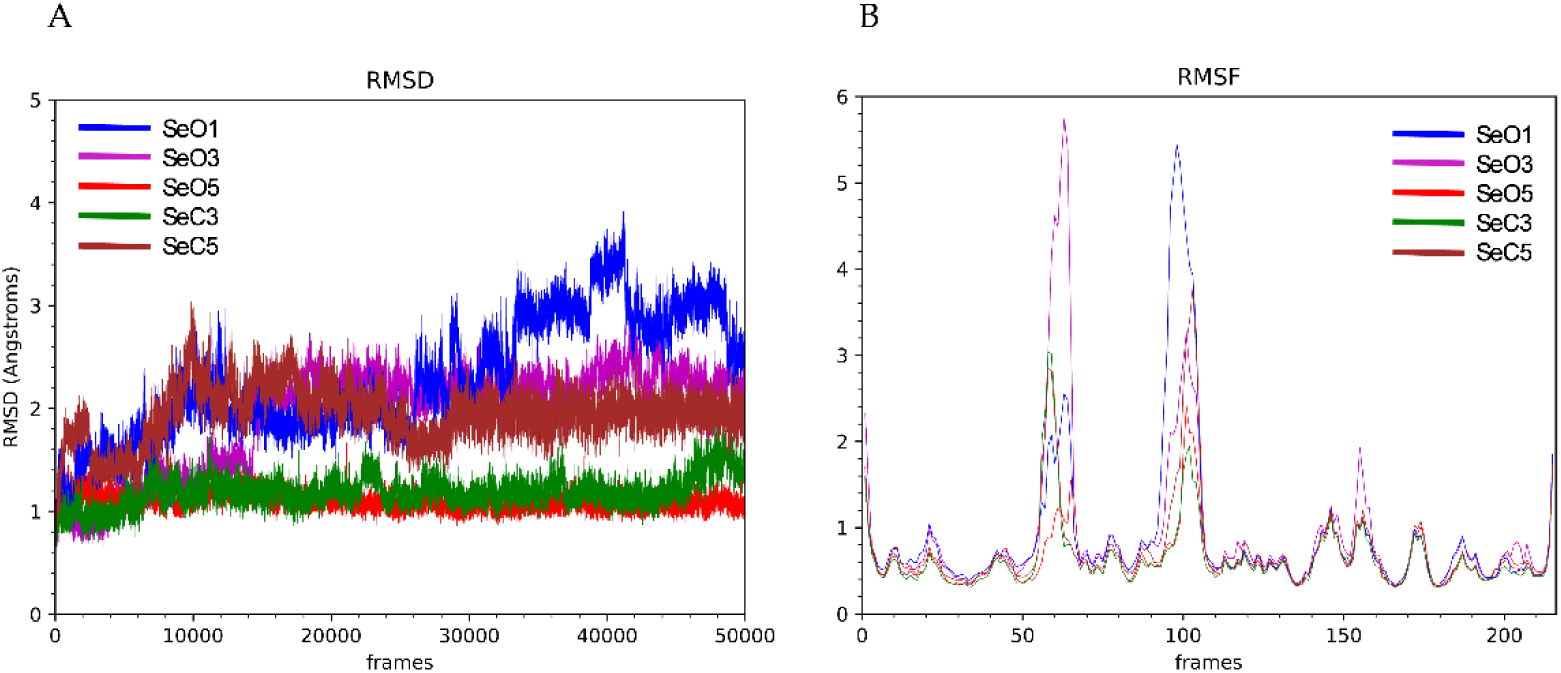
(A) Plot of the evolution of RMSD values during MD simulations of *Tbr*CATL complexed with the ligands ***Se*O1** (blue), ***Se*O3** (purple), ***Se*O5** (red), ***Se*C3** (green) and ***Se*C5** (brown). (B) RMSF plot of protein residues in complex with the ligands ***Se*O1** (blue), ***Se*O3** (purple), ***Se*O5** (red), ***Se*C3** (green) and ***Se*C5** (brown).

The solvent-accessible surface area (SASA) study suggested that the binding of the five ligands reduced the solvent-accessible surface area of the unbound *Tbr*CATL (**Table 3**). Also, the SASA graphs exhibited minimal fluctuations throughout the simulation trajectories (**Figure 4**). These findings align with the molecular mechanics Poisson-Boltzmann surface area (MMPBSA) binding free energy calculations, which yielded consistently negative values for all complexes (**Table 3**), thus, confirming thermodynamically favorable interactions between *Tbr*CATL and the *Se*-based ligands.

**Figure 4.**
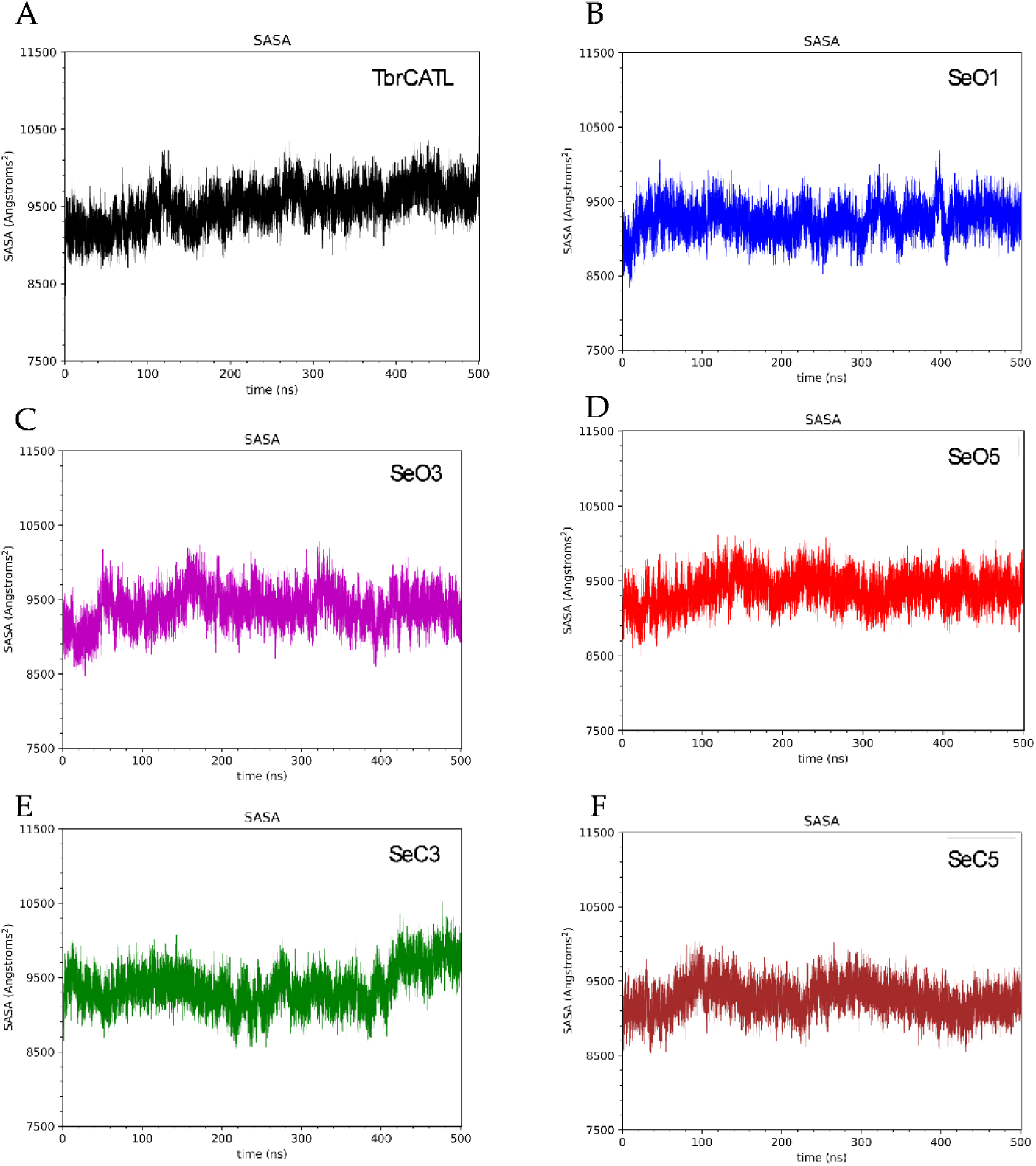
Solvent accessible surface area (SASA) of unbound *Tbr*CATL (A) and *Tbr*CATL complexed with ***Se*O1** (B), ***Se*O3** (C), ***Se*O5** (D), ***Se*C3** (E) and ***Se*C5** (F).

**Table 3.**
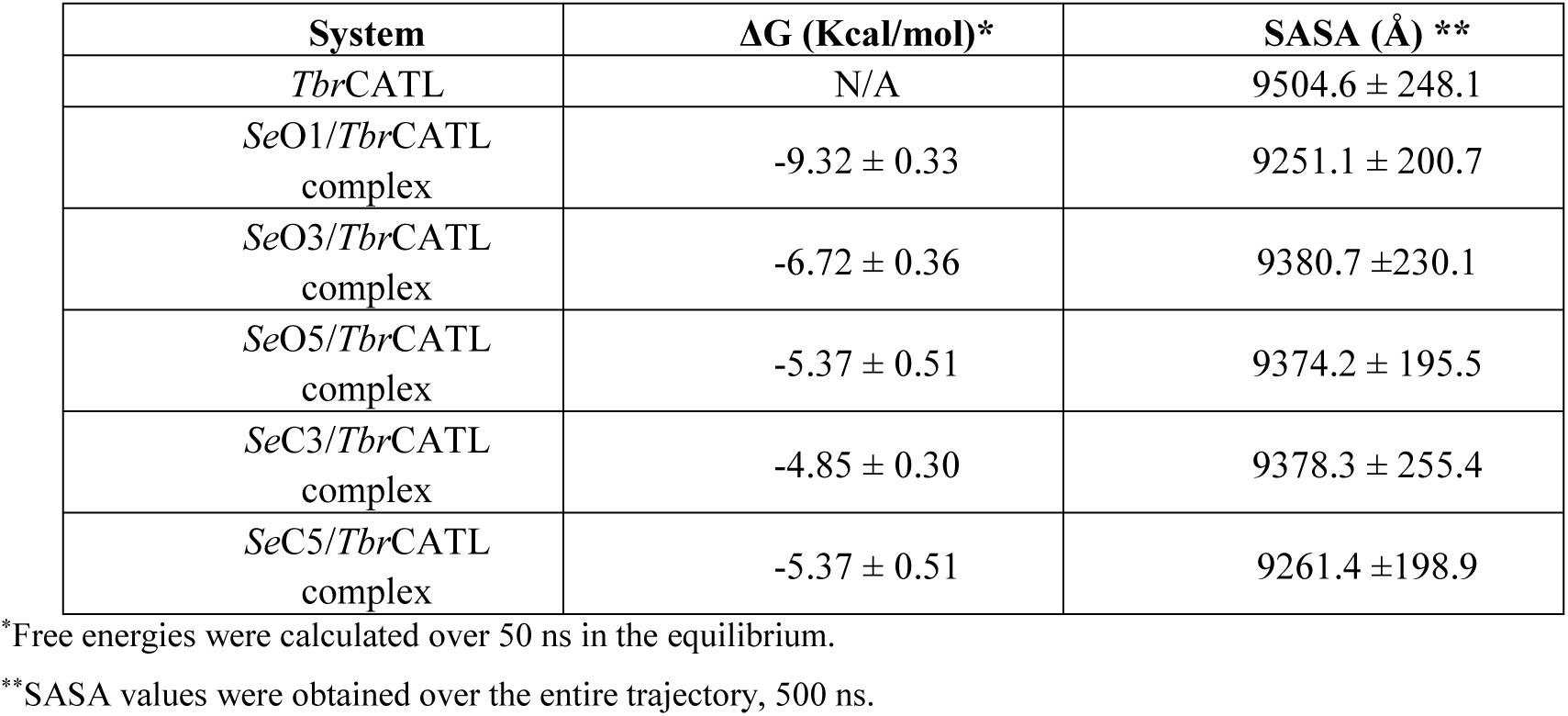
Predicted binding free energy values and mean SASA values obtained by MD simulations.

The interaction map of the open ligands ***Se*O1**, ***Se*O3**, and ***Se*O5** with *Tbr*CATL is shown in **Figure 5**. Compound ***Se*O1** forms hydrogen bonds through the selenosemicarbazone group hydrogens. Specifically, H3 interacts with the carbonyl oxygen of Asp^161^ and Gly^66^, while H1 forms polar contacts with Gly^66^ and H3 with Gly^62^ (see **Figure 5D** for ligand atom identification). Additionally, CH-π interactions are observed between the side chains of Leu^67^ and Ala^138^ at the S2 pocket and the aromatic ring of the ligand. A similar interaction pattern is displayed by ***Se*O3**, which also forms polar contacts through its H1 and H3 with Asp^161^ and Gly^66^, as well as CH-π interactions via its bromophenyl group with Leu^67^ and Ala^138^. In contrast, ***Se*O5** does not form any hydrogen bond with the protein. However, it still exhibits CH-π interactions through its aromatic ring and additional π-hole interactions between the electron-deficient nitrogen of the nitro group and the carbonyl groups of Gly^66^ and His^162^ (See Supporting **Table S2** for average distances of these interactions along with their occurrence during the simulation time).

**Figure 5.**
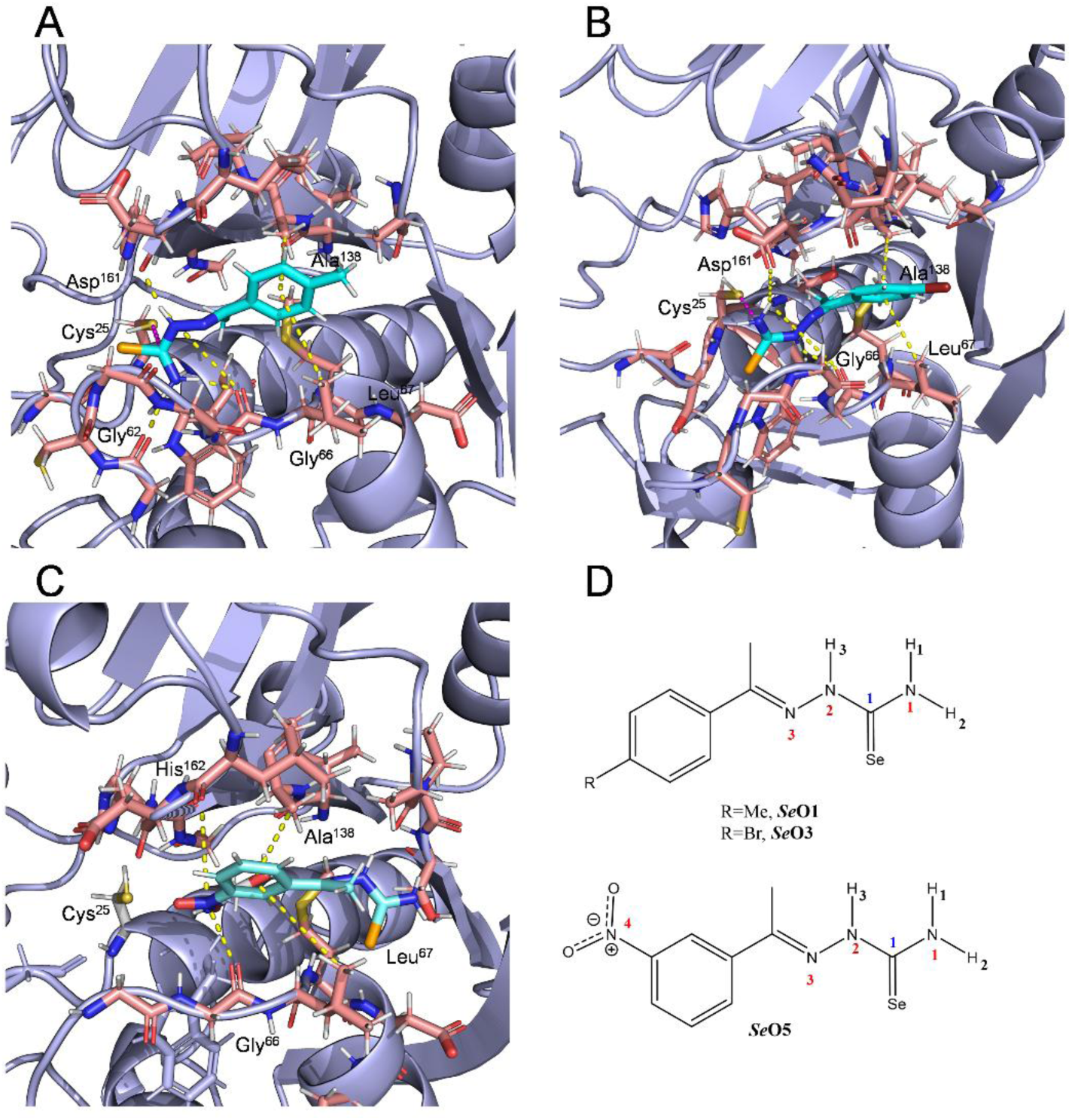
Representative frames of the binding modes for ***Se*O1** (A), ***Se*O3** (B) and ***Se*O5** (C) in the *Tbr*CATL active site (PDB: 2P7U) obtained from MD simulations. The frames selected here show the predominant conformations of the ligand-protein complex throughout the simulation. Yellow dashed lines indicate all interactions between the ligands and the protein, including hydrogen bonds, CH-π interactions and π-hole interactions. Pink dashed lines highlight the potential covalent bond formed between the thiol group of Cys^25^ and the selenocarbonyl group of the ligands. (D) Ligand atom labels involved in the recognition process.

### Mechanism of action: Radical scavenging capacity – antioxidant activity

As a possible alternative or additional mechanism of action, we evaluated the antioxidant capacity of the compounds shown in **Tables 1** and **2** at three different concentrations (0.06, 0.03 and 0.015 mg/ml), using the DPPH assay [49]. The assay was designed to measure the ability of the compounds to act as free radical scavengers or hydrogen donors. Quantitative data are shown in Supporting **Table S3,** and the graphical representation of the maximal antioxidant activity after 2 h is shown in **Figure 7**.

**Figure 6.**
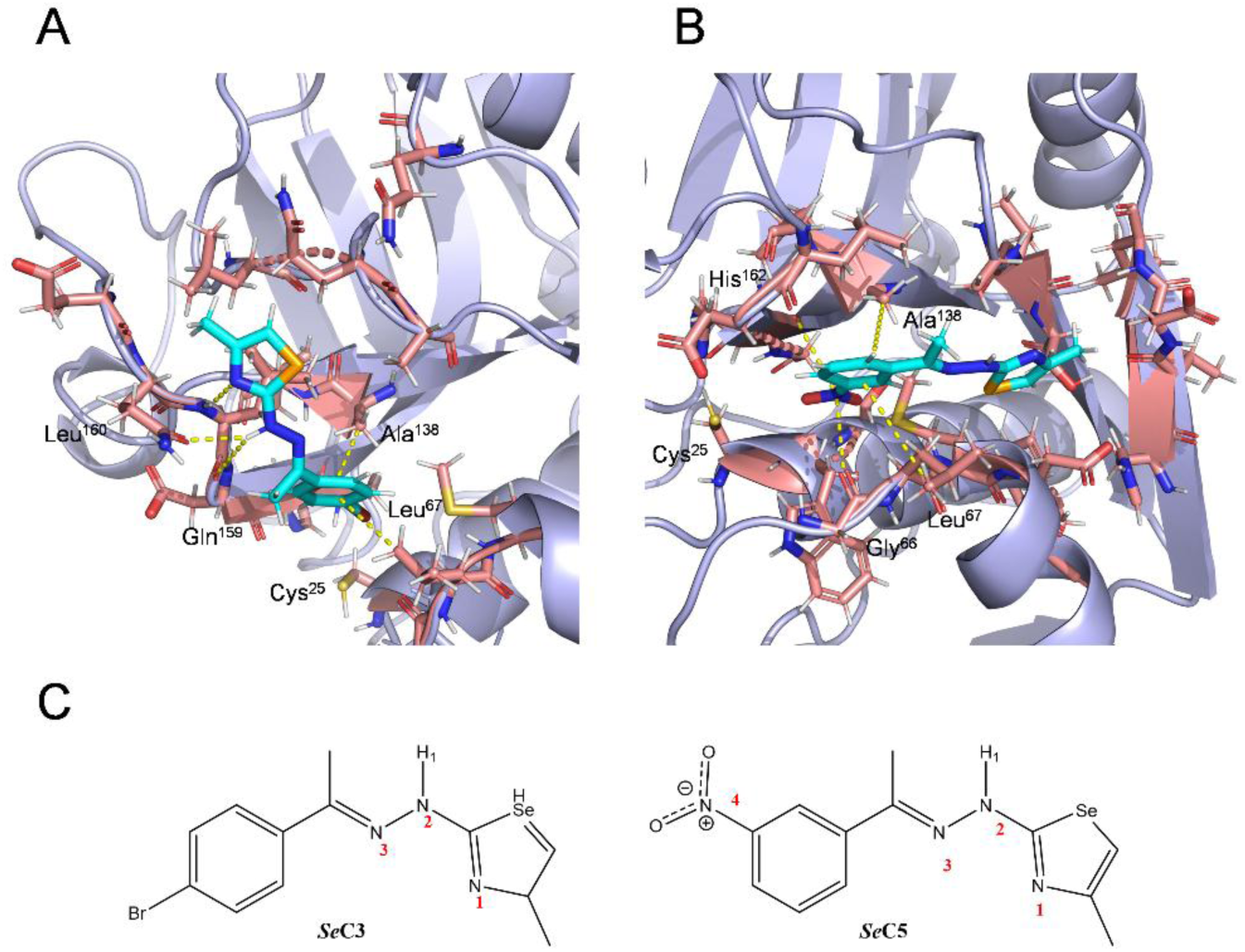
Representative frames of the binding modes of compounds ***Se*C3** (A) and ***Se*C5** (B) at the *Tbr*CATL active site (PDB: 2P7U), obtained from MD simulations. The frames selected here show the predominant conformations of the ligand-protein complex throughout the simulation. Yellow dashed lines indicate all interactions between the ligands and the protein, including hydrogen bonds, CH-π interactions and π-hole interactions. (C) Ligand atoms labels involved in the recognition process.

**Figure 7.**
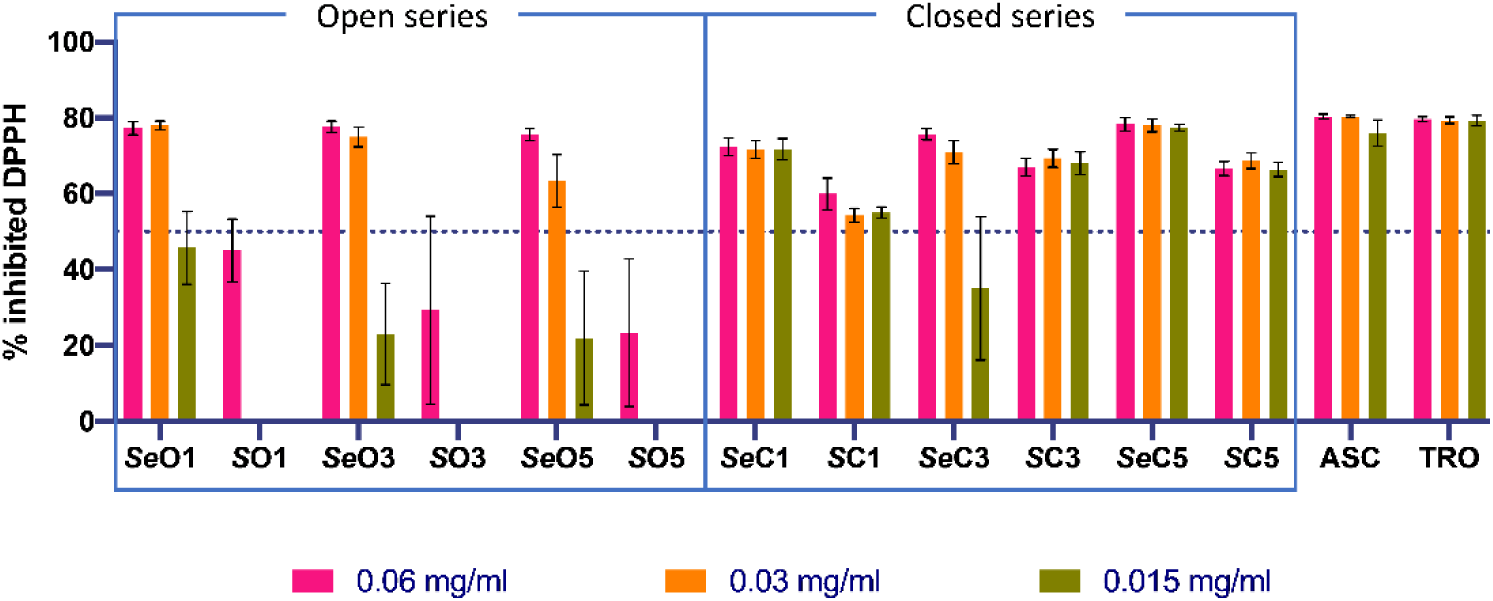
Antioxidant activity of the compounds of interest at the three tested concentrations after 2 h. ASC, ascorbic acid and TRO, trolox, were used as positive controls.

All compounds, except ***S*O1**, ***S*O3** and ***S*O5**, scavenged free radicals from 1,1-diphenyl-2-picrylhydrazyl (DPPH) by >50% after 2 h at the highest concentration tested (0.06 mg/ml). Therefore, these three compounds were not tested further (**Table S3**). Compared to the chalcogen counterparts, the *Se* compounds demonstrated enhanced antioxidant activity (70-80%) at 0.06 mg/ml and 0.03 mg/ml relative to their *S* analogues (40-70%), especially the open series, for which the *S* compounds failed to reach the 50% free radical inhibition threshold. This improved antioxidant activity by the *Se* compounds may be attributed to the higher reactivity of *Se*. In contrast, for the cyclic derivatives, the structure itself may enhance antioxidant activity, suggesting that the cyclic configuration plays a role alongside the chalcogen element. In this context, ***Se*C1** and ***Se*C5**, maintained similar inhibition values at the three concentrations tested (**Figure 7**). Indeed, ***Se*C5** is outstanding for its remarkable activity, which was comparable to that of the positive controls, ascorbic acid (ASC) and trolox (TRO).

### *In silico* prediction of ADME and drug-likeness properties

To predict their ADME and drug-likeness properties, all 44 compounds, and the positive drug controls, pentamidine and bortezomib, were analyzed using the SwissADME platform (http://www.swissadme.ch/) [50]. Data regarding molecular weight, lipophilicity, solubility, drug elimination and drug likeness criteria are available in the Supporting **Table S4**.

None of the 12 top hits (**Tables 1** and **2**) violated any of the drug-likeness criteria (Lipinski, Ghose, Veber and Egan rules) available on SwissADME platform (**Table S4**). In addition, all hits showed adequate oral bioavailability, scarce inhibition of cytochrome P450 (**Table S4**) and good gastrointestinal absorption (HIA; **Figure 8**). Also, we would note that that the cyclic derivatives, ***Se*C1**, ***Se*C3**, ***S*C1** and ***S*C3**, are more likely to cross the BBB than their open analogues (**Figure 8**). Data regarding passive gastrointestinal absorption (HIA) and brain access (BBB) are shown in the BOILED-Egg graph [51] (**Figure 8**). From this graph, we can also ascertain that only bortezomib (shaded in blue) is a substrate for P-glycoprotein (PGP+), a key protein responsible for efflux through biological membranes, giving us an idea of the mechanism of permeability used by our compounds.

**Figure 8.**
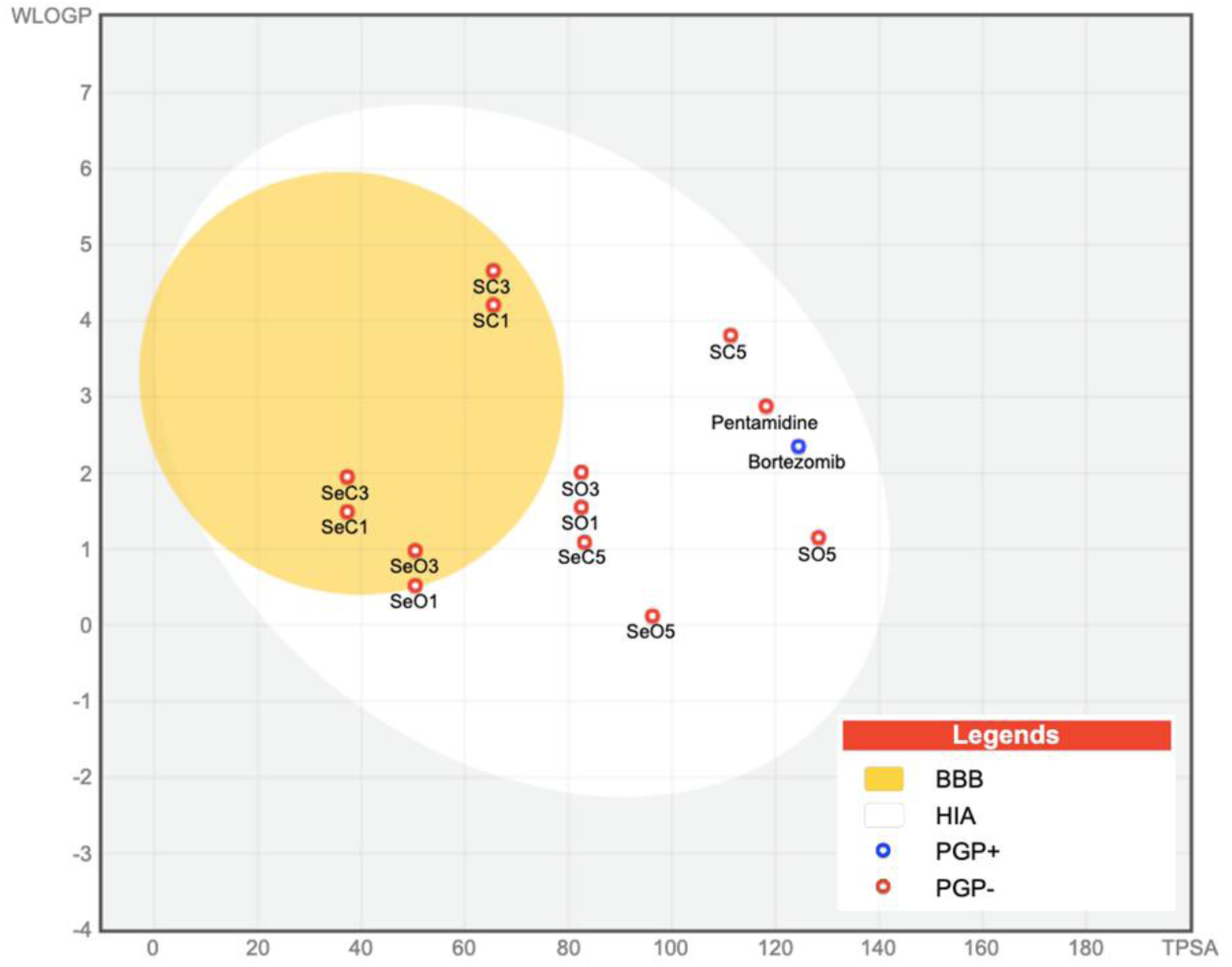
BOILED-Egg model of the 12 top hits and positive drug controls (pentamidine and bortezomib). The white region contains those compounds that are prone to be passively absorbed from the gastrointestinal tract (HIA). The yellow region (yolk) contains the molecules predicted to cross the BBB. Compounds colored in blue are predicted to be P-glycoprotein substrates (PGP+), whereas those in red are not (PGP-).

## DISCUSSION

Although *Se* has been neglected in medicinal chemistry, mainly because of its toxicity [52] and the comparison of *S-* and *Se-*containing compounds has been scarce. Accordingly, we implemented an isosteric replacement strategy for semicarbazones by changing *S* to *Se*. *S-*semicarbazones are the starting point for this work because they are established inhibitors of the trypanosomal cysteine proteases, Cz and *Tbr*CATL [28–30]. We also synthesized thiazoles and selenazoles to include structural variability and, for the first time, test selenazoles against *T. brucei*. Of 44 novel chalcogen derivatives tested against *T. brucei*, the *Se* derivatives were the most active (>70%). Structurally, nine of the active compounds (***Se*O1**, ***Se*O2**, ***Se*O3**, ***Se*O4**, ***Se*O5**, ***Se*O6**, ***Se*O9**, ***Se*O10**, ***Se*O11**) belong to the open series, whereas the remaining three (***Se*C1**, ***Se*C3**, ***Se*C5**) are closed derivatives. Comparing the EC_50_ values of these *Se-*derivatives, the open compounds (***Se*O3** and ***Se*O1**) are at least ten times better than their corresponding cyclic analogues (***Se*C3** and ***Se*C1**). Overall, the combination of the open conformation and *Se* improves activity against *T. brucei*, as demonstrated by the best compound, ***Se*O3** and its EC_50_ value of 0.47 ± 0.02 µM.

Regarding toxicity, 19 of the 22 *Se-*compounds were toxic, in contrast to five *S*-compounds. All of the trypanocidal *Se*-derivatives were cytotoxic to both HEK293 and HepG2 cells with similar CC_50_ values and SIs (**Table S1**). We only performed DR analysis for those compounds that were active against *T. brucei.* Of these, ***Se*O3** had the best SI_HEK293_ value of 6. Nevertheless, we found that the cyclic *Se-*derivatives were less toxic than their open counterparts. This is in accordance with other studies in the literature which report that thiazoles decrease toxicity [53–56]. This suggests that the structural configuration of the cyclic derivatives plays a key role in mitigating the inherent toxicity of *Se*, potentially due to increased stability. This finding is also consistent with our previous data published for C2C12 cells (immortalized mouse myoblasts) [40], and it will be necessary to design less toxic derivatives.

Related to the possible mechanism of action, as we reported for the main cysteine protease of *T. cruzi,* Cz [40], the open compounds are the more potent inhibitors of *Tbr*CATL (**Figure 2**). This is independent of the chalcogen compound present in the molecule, as the IC_50_ values between *S*- and *Se*-derivatives are very similar (0.30 to 17.90 nM). In addition, the nitro group in the aromatic ring may influence the interaction with *Tbr*CATL because all derivatives that contain this group (***Se*O5**, ***Se*C5**, ***S*O5** and ***S*C5**) inhibit the protease. For these reasons, we decided to investigate the interactions between the ligands and *Tbr*CATL utilizing MD simulations.

We performed MD simulations to determine the binding mode of ***Se*O1**, ***Se*O3**, ***Se*O5**, ***Se*C3** and ***Se*C5** with *Tbr*CATL. This involved studying the interactions between the protease and the ligands, and analyzing the potential formation of a covalent bond between the selenosemicarbazone moiety of open derivatives and the catalytic Cys^25^, as previously described [26,30]. The interaction maps for ***Se*O1** (**Figure 5A**) and ***Se*O3** (**Figure 5B**) indicates a common mode of recognition and binding to the *Tbr*CATL active site. The aromatic ring of both compounds participates in CH-π interactions and adopts a conformation whereby the selenosemicarbazone moiety forms multiple hydrogen bonds with various residues in the binding site. This ligand disposition satisfies the optimal geometric parameters necessary to potentially enable a nucleophilic attack by Cys^25^ on the selenocarbonyl group, leading to the formation of a covalent bond. This recognition mode is similar to that exhibited by structurally related *S*-semicarbazones previously described [30]. Of note, although the bromine-substituted closed derivative (***Se*C3**) displays the same pattern of CH-π interactions via its aromatic ring (**Figure 6A**), the selenazole counterpart points outward from the binding site, forming a different network of hydrogen bonds. Both polar and non-polar interactions are present in lower occurrences during the trajectory of the molecular dynamics (**Table S2**). Differences in contacts and the potential for covalent bond formation could explain the greater enzyme inhibition observed for the open compounds compared to their closed analogues (***Se*O3** *vs. **Se*****C3**). In addition to this information, the RMSD study of the ***Se*O1** and ***Se*O3** complexes shows greater deviation during the trajectory, likely due to the orientation of the selenosemicarbazone moiety towards the catalytic Cys^25^. This orientation may result in a decreased stabilization of the protein in both complexes, although it potentially facilitates the formation of a covalent bond.

The analysis of the interactions of ***Se*O5** (**Figure 5C**) and ***Se*C5** (**Figure 6B**) with *Tbr*CATL indicates that the nitro group of the nitrophenyl substituent plays a crucial role in enzyme inhibition. The electron-deficient nitrogen of the nitro aromatic group participates in π-hole interactions [57]. These polar interactions place the aromatic ring of the ligands in the same region of the binding site as the phenyl group of ***Se*O1** and ***Se*O3**, establishing CH-π interactions (**Figure 9**). The interactions involving the nitro group forces similar conformations for the open and closed chains of both ligands. The selenosemicarbazone chain of ***Se*O5** adopts a conformation that positions the selenocarbonyl group away from Cys^25^, preventing both covalent bond formation and polar interactions between the selenosemicarbazone hydrogens and the enzyme. The orientation of the chain containing the selenazole of ***Se*C5** matches the arrangement found in ***Se*O5**, underscoring the role of the nitro group in the binding mode of both ligands. This configuration of the nitro-aromatic derivatives aligns with our previous findings [40], in which interactions between ***Se*C5** and Cz were analyzed. Predictions for ***Se*C5** show that it interacts with Cz through a similar π-hole pattern as found for *Tbr*CATL, which is expected due to the structural similarities between both cysteine proteases [18,21,26]. These data align with the enzyme inhibition data, which show similar IC_50_ values for the open and closed derivatives bearing the nitro substituent (**Table 2**). Collectively, these findings provide valuable structural insights for the future design of compounds that effectively inhibit both *Tbr*CATL and Cz.

**Figure 9.**
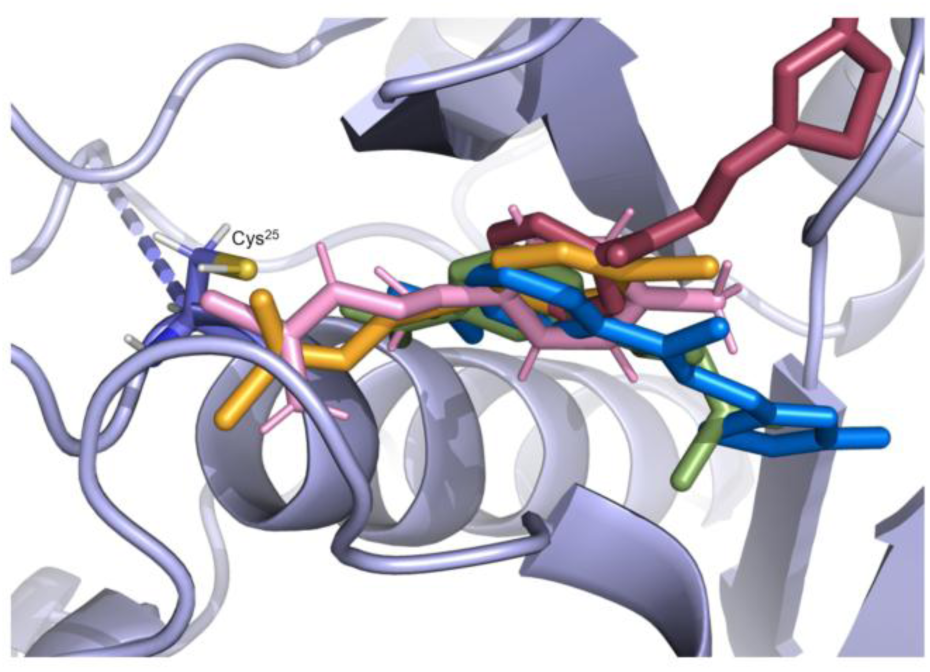
Superposition of the structures of ***Se*O1** (pink), ***Se*O3** (orange), ***Se*O5** (green), ***Se*C3** (red) and ***Se*C5** (blue) in the *Tbr*CATL active site (PDB 2P7U) obtained from MD simulations. The frames selected here show the predominant conformation of each ligand bound to the protein throughout the simulation.

In addition to the possible mechanism of *Tbr*CATL inhibition, the data generated for the trypanocidal activity suggest the presence of an additional mechanism(s) of action. This hypothesis is supported by the observation that the most potent trypanocide, ***Se*O3**, inhibits *Tbr*CATL (1.22 nM), whereas ***Se*C1** is active against *T. brucei*, but does not achieve >85% inhibition of the protease. This may be due to differences between the cell environment and the inhibition assay conditions. Accordingly, with the compounds from the groups shown in **Table 1** and **2**, we performed the DPPH assay to measure antioxidant activity, in the knowledge that *Se* is known for its free radical-scavenging activity [35,37]. The data reveal that the main contributor to the antioxidant activity measured is the structure. Specifically, all cyclic compounds were antioxidants, whereas only the *Se*-compounds from the open series acted as such (**Figure 7**). This suggests a direct correlation between the structural configuration and the stability of *Se* within the compounds such that the greater stability of the cyclic structures facilitates sustained antioxidant activity.

We consider that both possible mechanisms of action are important when designing future trypanocidal *Se*-containing compounds. This notion is reinforced by the fact that the derivatives, ***Se*O1**, ***Se*O3**, ***Se*O5**, ***Se*C3** and ***Se*C5** inhibit *Tbr*CATL between 0.30-17.90 nM (**Table 2**) and possess > 70% antioxidant activity at 0.06 mg/ml and 0.03 mg/ml (**Figure 7**). If one of these conditions is missing, the compound does not affect the parasite, *e.g*., ***Se*C1**, which shows antioxidant activity but does not inhibit *Tbr*CATL.

Last, based on *in silico* predictions of ADME and drug-likeness, all of the compounds tested are suitable for oral administration and meet all of the criteria for drug-likeness. Both the open and cyclic derivatives show good gastrointestinal absorption; however, the cyclic compounds are predicted to have a better likelihood of crossing the BBB compared to the open derivatives.

## CONCLUSIONS

In consideration of new pharmacological therapies for HAT, we evaluated 44 compounds against *T. brucei*. Using *S*-semicarbazones as a starting point, we employed a *S* to *Se* isosteric replacement strategy and divided compounds into two series of analogues, one containing open compounds and the other with the corresponding cyclic derivatives. Our main goal was to identify compounds that kill *T. brucei* and inhibit the validated drug target, *Tbr*CATL. Notably, *Se*-containing derivatives (open and closed) are more potent than their *S*-containing counterparts. The lead compound, ***Se*O3**, displayed low micromolar trypanocidal activity, low nanomolar inhibition of *Tbr*CATL, promising antioxidant activity, as well as predicted ADME and drug-likeness properties; however, its toxicity needs to be improved.

Regarding structural modifications, the selenazole derivatives demonstrated antioxidant activity, and, importantly, decreased *Se*-associated toxicity. Apart from the advantages of selenazole, further structural modifications should be performed in the *Se*-cyclic derivatives to improve both anti-parasitic activity and inhibition of *Tbr*CATL.

Related to the mechanism of action, we found that the combination of *Tbr*CATL inhibition and free radical-scavenging activity leads to *Se*-semicarbazone derivatives that kill *T. brucei*, as was the case for the lead compound, ***Se*O3**.

## METHODS

### Chemistry

The synthesis and characterization of the 44 compounds evaluated in this work were previously described by our group [40]. Their characterization included melting points, IR, ^1^H and ^13^C spectra. ^77^Se NMR was performed when possible. The purity (≥95%) of the compounds was determined by qNMR.

### *In vitro* antitrypanosomal assay

Bloodstream forms of *T. b. brucei* Lister 427 were grown in HMI-9 modified medium [58] supplemented with 20% heat-inactivated fetal bovine serum (FBS) in T25 suspension cell culture flasks at 37°C in 5% CO_2_ [59]. Trypanosomes were maintained in exponential growth phase and passaged every 48–72 h. Growth inhibition of *T. b. brucei* was determined using the SYBR Green cell viability assay [45]. Test compounds were diluted in DMSO and added to 96-well polystyrene assay plates to give final assay concentration of 10 μM (1 μL; 0.5% total DMSO). Fresh HMI-9 medium (99 μL/well) was added to the assay plate. Parasites in exponential phase were suspended at 2×10^5^ parasites/mL in HMI-9 medium and added to each well (100 μL) to a total density of 2×10^4^ trypanosomes/well. Assay plates were incubated at 37°C and 5% CO_2_ for 72 h, followed by addition of 50 µL/well of lysis solution (30 mM Tris, pH 7.5, 0.012% saponin, 0.12% Triton X-100 and 7.5 mM ETDA) containing 0.3 µL/mL SYBR Green I (10,000X in DMSO; Invitrogen, Carlsbad, CA). Assay plates were incubated in the dark for 1 h at room temperature. Fluorescence was measured at 485 nm and 535 nm excitation and emission wavelengths, respectively, using a 2104 EnVision® multilabel plate reader. The viability of each well was normalized to positive and negative controls in each assay plate (pentamidine was used as the positive control). The screen was performed in technical quadruplicate. DR curves were generated for compounds inhibiting parasite growth by ≥ 70%. Eight-point concentration-response curves were prepared and EC_50_ values were calculated with GraphPad Prism, version 9.3 (San Diego, CA) using a sigmoidal four parameter logistic curve. EC_50_ data were generated from three independent experiments each performed in duplicate.

### *In vitro* cytotoxicity evaluation

The HEK293 and HepG2 cells were cultured in DMEM supplemented with 10% heat-inactivated FBS and 1% penicillin-streptomycin [60,61]. Cells were grown in T75 cell culture flasks maintained at 37°C in 5% CO_2_ and sub-cultured when at 60–80% cell confluence. Cytotoxicity was measured using the Promega CellTiter-Glo reagent (G7572) [60,61]. Test compounds were diluted in DMSO and added to 96-well polystyrene assay plates to give a final assay concentration of 10 μM. Fresh medium was added to the assay plate (49 μL/well). HEK293 or HepG2 cells were suspended to 1×10^5^ cells/mL in DMEM and added to each well (50 μL) for a total density of 5×10^3^ cells/well. Assay plates were incubated at 37°C and 5% CO_2_ for 48 h, followed by addition of 25 μL/well of CellTiter-Glo. Luminescence was measured using a 2104 EnVision® multilabel plate reader. The viability of each well was normalized to positive and negative controls in each assay plate (bortezomib was used as the positive control). CC_50_ values were calculated with GraphPad Prism, version 9.3 (San Diego, CA) using a sigmoidal four parameter logistic curve. CC_50_ data were generated from three independent experiments each performed in duplicate.

### Enzyme inhibition assays

All experiments were performed in a 384-well black microplate, using a Synergy HTX (Biotek) plate reader at 25°C with absorption/emission wavelengths of 360/460 nm. Assays were performed in triplicate wells and DMSO was used as a vehicle control. Enzyme activity was measured and normalized to DMSO controls, and the inhibitor E-64 (10 µM) was used as a positive control for all assays. Experiments were performed in triplicate, with at least two independent assays (n = 6 data points). IC_50_ values were determined through nonlinear regression. The data obtained were analyzed using GraphPad Prism 9.0 (GraphPad Software, La Jolla, California, US).

### *Tbr*CATL inhibition assays

The screening assay was performed using 0.5 nM of *Tbr*CATL diluted in 0.1 M sodium acetate, 1 mM dithiothreitol (DTT), and 0.01% Triton X-100, pH 5.5. *Tbr*CATL was expressed and purified as described [62]. Enzyme the compound (10 µM), were incubated for 10 min at room temperature. Then the substrate, Z-Phe-Arg-amidomethylcoumarin (Z-FR-AMC; Sigma-Aldrich, C9521), was added (30 µL diluted in the same assay buffer) to yield a final concentration of 2.5 µM and the assay allowed to proceed for 10 min. The IC_50_ values of compounds inhibiting more than 85% at 10 µM and active against *T. brucei* (antiparasitic activity ≥ 70%) were calculated. For those compounds inhibiting enzyme activity by >85%, DR assays were performed over 15 two-fold serial dilutions starting at 0.5 or 20 µM.

### *h*CatL inhibition assays

Recombinant *h*CatL was purchased from R&D Systems (952-CY) and activated according to the manufacturers protocol. This enzyme, *h*CatL, is used to test compounds’ selectivity because it is the human homolog of *Tbr*CATL. The assay was modified from [63], using 25 pM of the enzyme diluted in 40 mM sodium acetate, 5 mM DTT, 100 mM NaCl, 1 mM EDTA and 0.01% Triton X-100, pH 5.5. Enzyme and compound (10 µM), were incubated for 30 min at room temperature. Then Z-FR-AMC was added (30 µL diluted in the same assay buffer) to yield a final concentration of 25 µM and the assay allowed to proceed for 10 min. DR assays were performed over 15 two-fold serial dilutions starting at 0.5 or 20 µM.

### Molecular dynamics

The structure deposited in the Protein Data Bank (PDB) under the ID 2P7U [48] prepared the starting structures to run the Molecular Dynamics Simulations (MD). The ligand of the 2P7U structure was replaced by the selenium derivatives ***SeO1****, **SeO3**, **SeO5**, **SeC3*** and ***SeC5*** before running the MD simulation. Both the systems preparation and the simulations were performed in the AMBER 18 suite software. The protocol for the systems preparation and the MD simulations is detailed as follows. Firstly, the system is neutralized by adding sodium ions and later immersed in a cubic box of 10 Å length, in each direction from the end of the protein, using TIP3P water parameters.

The force fields used to obtain topography and coordinates files were ff14SB [64] and GAFF [65]. The first step of the simulation protocol followed to run the MD simulations is a minimization of the solvent molecules position only, keeping the solute atom positions restrained, and the second stage minimizes all the atoms in the simulation cell. Heating the system is the third step, which gradually raises the temperature 0 to 300 K under a constant volume (ntp = 0) and periodic boundary conditions. In addition, Harmonic restraints of 10 kcal/mol^-1^ were applied to the solute, and the Berendsen temperature coupling scheme [66] was used to control and equalize the temperature. The time step was kept at 2 fs during the heating phase. Long-range electrostatic effects were modelled using the particle-mesh-Ewald method [67]. The Lennard-Jones interactions cut-off was set at 8 Å. An equilibration step for 100 ps with a 2fs time step at a constant pressure and temperature of 300 K was performed prior to the production stage. The trajectory production stage kept the equilibration step conditions and was prolonged for 500 ns longer at the 1 fs time step. In addition, the selenium derivative required a previous preparation step where the parameters and charges were generated by using the antechamber module of AMBER, using the GAFF force field and AM1-BCC method for charges [68].

The interaction free energy between the ligands and *Tbr*CATL was estimated using the Molecular Mechanics Poisson-Boltzmann Surface Area (MM/PBSA) method.

### Evaluation of antioxidant activity

The antioxidant capacity of the selected Se-compounds was tested using the DPPH assay as described [49]. DPPH forms a free radical that is stable at room temperature. If the tested compounds are antioxidant, they neutralize the free radicals to donating a proton [69]. The compounds were tested at three different concentrations (0.06, 0.03 and 0.015 mg/ml) and we used two positive controls: ascorbic acid (ASC) and trolox (TRO). A solution of DPPH (2,2-diphenyl-1-picrylhydrazyl) in methanol (0.04 mg/ml, preserved in the dark) was prepared and 100 µL of this stock solution was added to 100 µL of tested compounds solution. The color change, from purple (radical) to yellow (reduced), was measured at 517 nm at different time points (0, 5, 15, 30, 60, 90 and 120 min). The experiment was performed three times in triplicate and the measurements were recorded on a BioTek PowerWave XS spectrophotometer (Biotek). Data were collected using BioTek Gen5 Microplate reader and Imager software (Agilent, version 3.12). Data were generated from three independent experiments, each performed in triplicate (n=9), and expressed as a percentage of inhibited DPPH (mean ± standard error of the mean (SEM)) using the following formula:

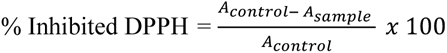

where A_control_ refers to the absorbance of the negative control and A_sample_ refers to the absorbance of the tested compounds.

### *In silico* prediction of ADME and drug-likeness properties

SwissADME platform (http://www.swissadme.ch/) (accessed on 15^th^ January 2025) was used to analyze the physicochemical and pharmacokinetic characteristics of the 44 compounds evaluated in this work. This platform is freely provided by the Swiss Institute of Bioinformatics (SIB) and provides information on bioavailability and ADME parameters (Absorption, Distribution, Metabolism and Elimination) [50].

## SUPPLEMENTARY MATERIALS

The following supporting information can be downloaded at: https://pubs.acs.org/doi/xxx Table S1: Data for the 44 compounds screened against *T. brucei brucei* Lister 427,HEK293 and HepG2 cells; Figure S1: DR of *S*-compounds against *Tbr*CATL (A) and *h*CatL (B); Table S2: Average main interaction distances between the ligands and the protein as complexes, and their occurrence during the MD simulation trajectories; Table S3: DPPH data for three compound concentrations after 2 h; Table S4: ADME and drug-likeness properties predicted for 44 compounds using SwissADME.

## ACKNOWLEDGMENTS

MRH acknowledges all of the personnel at the CDIPD, UC San Diego, and at UNAV who were involved in this project.

