## Supplementary material for "Chalcogen derivatives for the treatment of African trypanosomiasis: biological evaluation of thio and seleno- semicarbazones and their azole derivatives": Suppl data

**Contents:**

1. Biological data
2. Inhibition of *Tbr*CATL and *h*CatL by selected *S*-compounds
3. Molecular Dynamics data
4. DPPH data
5. *In silico* ADME and drug-likeness predictions

### 1. Biological data

**Table S1.** Data for the 44 compounds screened against *T. brucei brucei* Lister 427, HEK293 and HepG2 cells.

| Structure | Compound | *T. brucei* inhibition (%)[a] | HEK293 inhibition (%)[b] | HepG2 inhibition (%)[c] | $EC_{50}$ *T. brucei* (μM)[d] | $CC_{50}$ HEK293 (μM)[e] | $SI_{HEK293}$ [f] | $CC_{50}$ HepG2 (μM)[e] | $SI_{HepG2}$ [f] |
|---|---|---|---|---|---|---|---|---|---|
| 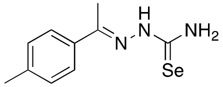 | ***Se*O1** | 100 ± 1 | 100 ± 1 | 80 ± 1 | 0.98 ± 0.12 | 4.27 ± 0.22 | 4 | 4.17 ± 0.60 | 4 |
| 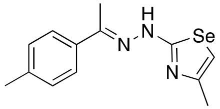 | ***Se*C1†** | 72 ± 6 | 77 ± 2 | 65 ± 9 | 9.80 ± 0.63 | 11.97 ± 4.94 | 1 | 10.98 ± 0.88 | 1 |
| 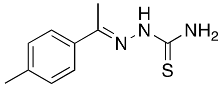 | **SO1** | 44 ± 7 | 30 ± 3 | 50 ± 4 | ND | ND | ND | ND | ND |
| 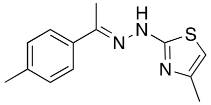 | **SC1†** | 30 ± 6 | 24 ± 7 | 4 ± 15 | ND | ND | ND | ND | ND |
| 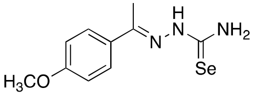 | ***Se*O2** | 100 ± 3 | 100 ± 2 | 78 ± 3 | 2.36 ± 0.02 | 5.81 ± 0.11 | 2 | 4.12 ± 0.44 | 2 |
| 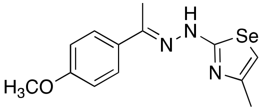 | ***Se*C2†** | 23 ± 7 | 80 ± 6 | 40 ± 5 | ND | ND | ND | ND | ND |
| 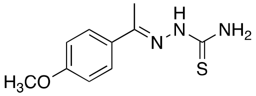 | **SO2** | 6 ± 2 | 22 ± 7 | 34 ± 9 | ND | ND | ND | ND | ND |

| | | | | | | | | | |
|---|---|---|---|---|---|---|---|---|---|
| 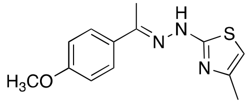 | **SC2†** | 27 ± 5 | 33 ± 8 | 6 ± 3 | ND | ND | ND | ND | ND |
| 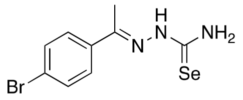 | ***Se*O3** | 100 ± 2 | 100 ± 0 | 71 ± 15 | 0.47 ± 0.02 | 2.82 ± 0.11 | 6 | 2.70 ± 0.16 | 6 |
| 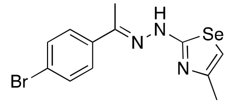 | ***Se*C3†** | 99 ± 2 | 88 ± 3 | 80 ± 2 | 6.04 ± 0.72 | 5.90 ± 1.22 | 1 | 4.68 ± 0.14 | 1 |
| 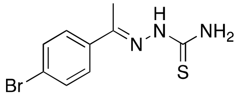 | **SO3** | 28 ± 4 | 76 ± 2 | 65 ± 4 | ND | ND | ND | ND | ND |
| 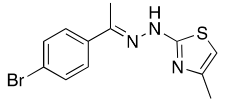 | **SC3†** | 38 ± 9 | 64 ± 2 | 46 ± 7 | ND | ND | ND | ND | ND |
| 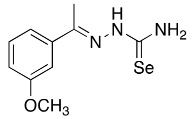 | ***Se*O4** | 100 ± 1 | 100 ± 2 | 74 ± 4 | 3.28 ± 0.29 | 3.25 ± 0.30 | 1 | 4.18 ± 0.25 | 1 |
| 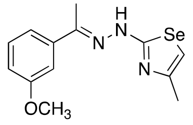 | ***Se*C4†** | 36 ± 4 | 60 ± 7 | 66 ± 5 | ND | ND | ND | ND | ND |
| 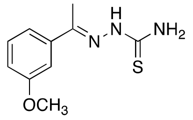 | **SO4** | 29 ± 5 | 31 ± 4 | 28 ± 6 | ND | ND | ND | ND | ND |
| 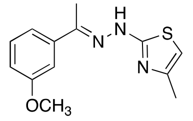 | **SC4†** | 54 ± 5 | 40 ± 4 | 52 ± 3 | ND | ND | ND | ND | ND |

| | | | | | | | | | |
|---|---|---|---|---|---|---|---|---|---|
| 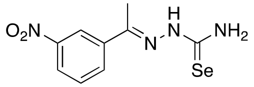 | ***Se*O5** | 100 ± 1 | 100 ± 0 | 85 ± 0 | 5.83 ± 0.72 | 2.29 ± 0.26 | 0 | 5.54 ± 1.47 | 1 |
| 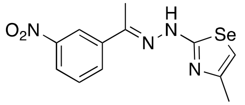 | ***Se*C5†** | 100 ± 2 | 81 ± 7 | 77 ± 6 | 10.53 ± 1.02 | 9.00 ± 1.36 | 1 | 11.27 ± 0.97 | 1 |
| 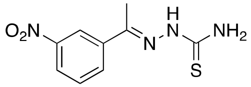 | **SO5** | 31 ± 5 | 14 ± 9 | 39 ± 9 | ND | ND | ND | ND | ND |
| 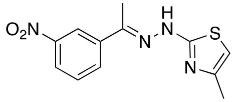 | **SC5†** | 40 ± 8 | 30 ± 3 | 17 ± 2 | ND | ND | ND | ND | ND |
| 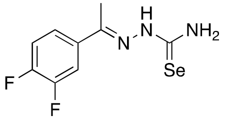 | ***Se*O6** | 100 ± 0 | 100 ± 0 | 75 ± 1 | 2.57 ± 0.15 | 3.69 ± 0.37 | 2 | 4.43 ± 0.11 | 2 |
| 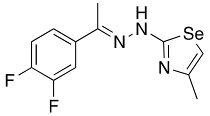 | ***Se*C6†** | 37 ± 4 | 48 ± 9 | 55 ± 5 | ND | ND | ND | ND | ND |
| 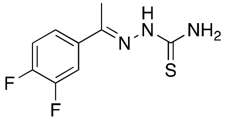 | **SO6** | 44 ± 4 | 36 ± 5 | 36 ± 1 | ND | ND | ND | ND | ND |
| 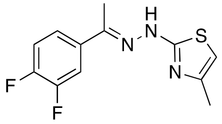 | **SC6†** | 22 ± 3 | 35 ± 1 | 0 ± 7 | ND | ND | ND | ND | ND |
| 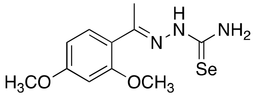 | ***Se*O7** | 12 ± 1 | 99 ± 2 | 2 ± 3 | ND | ND | ND | ND | ND |

| | | | | | | | | | |
|---|---|---|---|---|---|---|---|---|---|
| 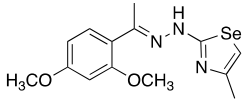 | ***Se*C7†** | 15 ± 4 | 38 ± 7 | 4 ± 6 | ND | ND | ND | ND | ND |
| 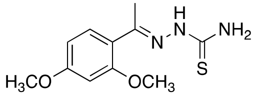 | **SO7** | 34 ± 5 | 13 ± 3 | 1 ± 1 | ND | ND | ND | ND | ND |
| 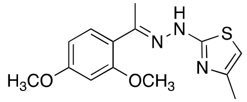 | **SC7†** | 39 ± 8 | 23 ± 1 | 0 ± 7 | ND | ND | ND | ND | ND |
| 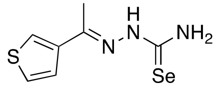 | ***Se*O8** | 11 ± 1 | 100 ± 1 | 62 ± 2 | ND | ND | ND | ND | ND |
| 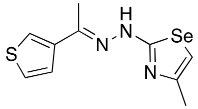 | ***Se*C8†** | 25 ± 6 | 37 ± 3 | 22 ± 3 | ND | ND | ND | ND | ND |
| 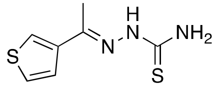 | **SO8** | 23 ± 2 | 13 ± 1 | 0 ± 6 | ND | ND | ND | ND | ND |
| 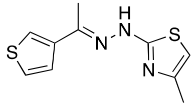 | **SC8†** | 0 ± 1 | 28 ± 1 | 0 ± 6 | ND | ND | ND | ND | ND |
| 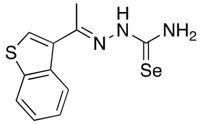 | ***Se*O9** | 100 ± 1 | 100 ± 0 | 67 ± 5 | 4.80 ± 0.33 | 3.46 ± 0.12 | 1 | 4.93 ± 0.41 | 1 |
| 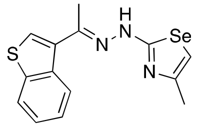 | ***Se*C9†** | 58 ± 1 | 73 ± 9 | 85 ± 5 | ND | ND | ND | ND | ND |

| | | | | | | | | | |
|---|---|---|---|---|---|---|---|---|---|
| 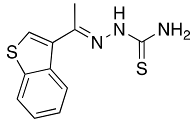 | **SO9** | 18 ± 6 | 58 ± 6 | 17 ± 5 | ND | ND | ND | ND | ND |
| 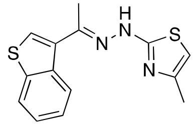 | **SC9†** | 39 ± 7 | 55 ± 7 | 9 ± 10 | ND | ND | ND | ND | ND |
| 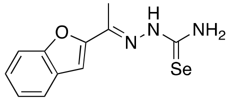 | ***Se*O10** | 100 ± 1 | 100 ± 1 | 82 ± 2 | 1.86 ± 0.33 | 3.00 ± 0.15 | 2 | 5.08 ± 0.27 | 3 |
| 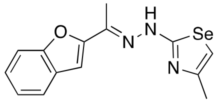 | ***Se*C10†** | 51 ± 7 | 65 ± 4 | 69 ± 10 | ND | ND | ND | ND | ND |
| 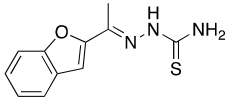 | **SO10** | 0 ± 8 | 28 ± 3 | 54 ± 5 | ND | ND | ND | ND | ND |
| 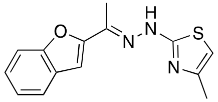 | **SC10†** | 46 ± 6 | 42 ± 0 | 49 ± 9 | ND | ND | ND | ND | ND |
| 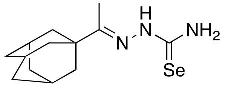 | ***Se*O11** | 100 ± 1 | 100 ± 4 | 66 ± 5 | 5.16 ± 1.06 | 10.50 ± 0.36 | 2 | 6.01 ± 0.69 | 1 |
| 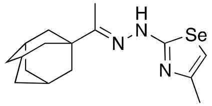 | ***Se*C11†** | 37 ± 0 | 64 ± 10 | 66 ± 2 | ND | ND | ND | ND | ND |
| 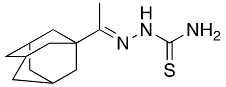 | **SO11** | 0 ± 4 | 75 ± 0 | 0 ± 12 | ND | ND | ND | ND | ND |

| | | | | | | | | | |
|---|---|---|---|---|---|---|---|---|---|
| 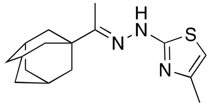 | **SC11†** | 19 ± 8 | 39 ± 3 | 0 ± 13 | ND | ND | ND | ND | ND |
| 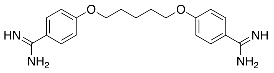 | **Pentamidine** | 100 ± 2 | NA | | 0.014 ± 0.002 | NA | | | |
| 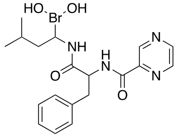 | **Bortezomib** | ND | 100 ± 2 | 100 ±1 | ND | 0.015 ± 0.002 | | 0.015 ± 0.003 | |

[a]*T. brucei* growth inhibition was calculated at 10 µM and is shown as a percentage (± SD).

[b]HEK293 growth inhibition was calculated at 10 µM and is shown as percentage (± SD).

[c]HepG2 growth inhibition was calculated at 10 µM and is shown as percentage (± SD).

[d]$EC_{50}$ values (mean ± SD) were calculated from three independent experiment each in duplicate.

[e]$CC_{50}$ values (mean ± SD) were calculated from three independent experiment each in duplicate.

[f]SI is the selectivity index of the compound, *i.e.*, $CC_{50}$ HEK293/$EC_{50}$ or $CC_{50}$ HepG2/$EC_{50}$.

ND: no data.

†These compounds were tested in their hydrochloride form.

### 2. Inhibition of *Tbr*CATL and *h*CatL by selected *S*-compounds

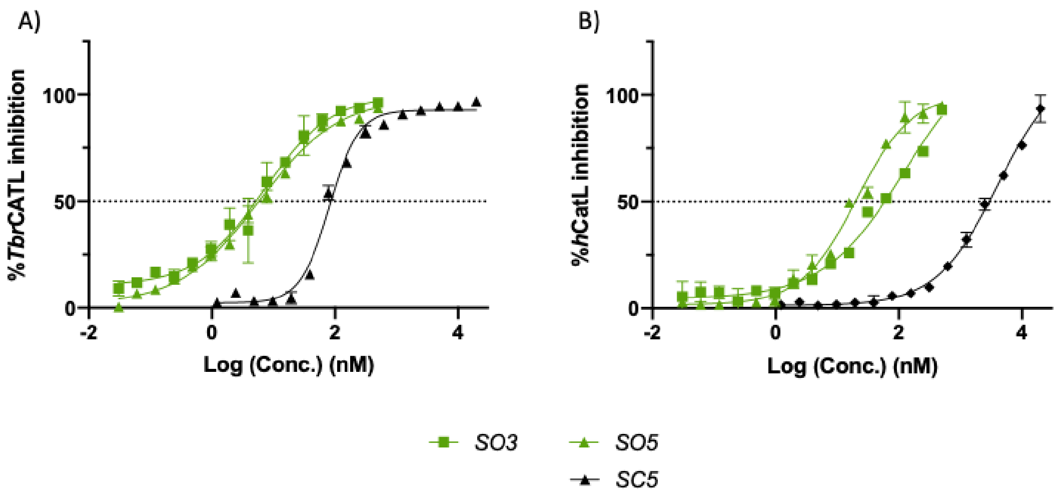


**Figure S1.** DR of *S*-compounds against *Tbr*CATL (A) and *h*CatL (B).

### 3. Molecular Dynamics data

**Table S2.** Average main interaction distances between the ligands and the protein as complexes, and their occurrence during the MD simulation trajectories.

| Ligand | Type of interaction | Involved Atoms | Distance (Å) | Occurrence (%) |
|---|---|---|---|---|
| ***Se*O1** | H bond | Asp 161 (O)- Ligand N2 (H3) | 2.8 | 49 |
| | H bond | Gly 66 (O)-Ligand N2 (H3) | 2.8 | 14 |
| | H bond | Gly 66 (O)-Ligand N1 (H1) | 2.9 | 26 |
| | H bond | Gly 62 (O)-Ligand N1 (H2) | 2.9 | 36 |
| | CH-π | Leu 67 (Cδ2)- phenyl ring | 3.7 | 48 |
| | CH-π | Ala 138 (Cβ)- phenyl ring | 3.6 | 5 |
| ***Se*O3** | H bond | Asp 161 (O)- Ligand N1 (H1) | 2.8 | 10 |
| | H bond | Gly 66 (O)-Ligand N2 (H3) | 2.9 | 10 |
| | H bond | Gly 66 (O)-Ligand N1 (H1) | 2.8 | 5 |
| | CH-π | Leu 67 (Cδ2)- phenyl ring | 3.7 | 48 |
| | CH-π | Ala 138 (Cβ)- phenyl ring | 3.7 | 51 |
| ***Se*O5** | CH-π | Leu67 (Cδ2)- phenyl ring | 3.8 | 10 |
| | CH-π | Ala 138 (Cβ)- phenyl ring | 3.8 | 19 |
| | Π-Hole | Gly66 (O)-Ligand N4 | 3.2 | 11 |
| | Π-Hole | His162(O)-Ligand N4 | 3.4 | 2 |
| ***Se*C3** | H bond | Leu 160 (NH)-Ligand N1 | 2.9 | 5 |
| | H bond | Leu 160 (O)-Ligand N2(H) | 2.9 | 5 |
| | H bond | Gln 159 (Oε)-Ligand N2(H) | 2.9 | 1 |
| | CH-π | Leu67 (Cδ2)- phenyl ring | 3.7 | 15 |
| | CH-π | Ala 138 (Cβ)- phenyl ring | 3.7 | 18 |
| ***Se*C5** | CH-π | Leu67 (Cδ2)- phenyl ring | 3.7 | 24 |
| | CH-π | Ala 138 (Cβ)- phenyl ring | 3.8 | 13 |
| | Π-Hole | Gly66 (O)-Ligand N4 | 3.2 | 29 |
| | Π-Hole | His162(O)-Ligand N4 | 3.3 | 5.2 |

### 4. DPPH data

**Table S3.** DPPH data for three compound concentrations after 2 h.

| Compound | 0.06 mg/ml* | 0.03 mg/ml* | 0.015 mg/ml* |
|---|---|---|---|
| ***Se*O1** | 77.33 ± 1.76 | 78.00 ± 1.16 | 45.67 ± 9.56 |
| ***Se*C1** | 72.33 ± 2.33 | 71.67 ± 2.40 | 71.67 ± 2.85 |
| ***S*O1** | 45.00 ± 8.33 | NT | NT |
| ***S*C1** | 60.00 ± 4.16 | 54.33 ± 1.86 | 55.00 ± 1.53 |
| ***Se*O3** | 77.67 ± 1.45 | 75.00 ± 2.65 | 23.00 ± 13.43 |
| ***Se*C3** | 75.67 ± 1.45 | 71.00 ± 3.06 | 35.00 ± 18.82 |
| ***S*O3** | 29.33 ± 24.77 | NT | NT |
| ***S*C3** | 67.00 ± 2.31 | 69.33 ± 2.33 | 68.00 ± 3.06 |
| ***Se*O5** | 75.67 ± 1.67 | 63.33 ± 6.89 | 22.00 ± 17.67 |
| ***Se*C5** | 78.33 ± 1.76 | 78.00 ± 1.73 | 77.33 ± 0.88 |
| ***S*O5** | 23.33 ± 19.46 | NT | NT |
| ***S*C5** | 66.67 ± 1.86 | 68.67 ± 2.03 | 66.33 ± 1.86 |
| **ASC$^{+}$** | 80.33 ± 0.67 | 80.33 ± 0.33 | 76.00 ± 3.52 |
| **TRO$^{+}$** | 79.67 ± 0.67 | 79.33 ± 0.882 | 79.33 ± 1.33 |

*mg/ml (expressed as the mean ± SEM of three independent experiments each performed in triplicate)

$^{+}$ASC: ascorbic acid; TRO: Trolox. Both compounds used as positive control.

NT: no tested.

### 5. *In silico* predictions *of* ADME and drug-likeness properties

**Table S4.** ADME and drug-likeness properties predicted for 44 compounds using SwissADME

| Compound | MW[a] | Log P[b] | Solubility[c] | Drug elimination parameters | | | | | Drug-likeness | | | |
|---|---|---|---|---|---|---|---|---|---|---|---|---|
| | | | | CYP1A2 inhibitor | CYP2C19 inhibitor | CYP2C9 inhibitor | CYP2D6 inhibitor | CYP3A4 inhibitor | Lipinski violations | Ghose violations | Veber violations | Egan violations |
| ***Se*O1** | 254.19 | 0.85 | Soluble | No | No | No | No | No | 0 | 0 | 0 | 0 |
| ***Se*C1** | 328.70 | 1.99 | Moderately soluble | No | No | No | No | No | 0 | 0 | 0 | 0 |
| ***S*O1** | 207.30 | 1.89 | Soluble | No | No | No | No | No | 0 | 0 | 0 | 0 |
| ***S*C1** | 281.80 | 3.16 | Moderately soluble | Yes | No | No | No | No | 0 | 0 | 0 | 0 |
| ***Se*O2** | 270.19 | 0.50 | Soluble | No | No | No | No | No | 0 | 0 | 0 | 0 |
| ***Se*C2** | 344.70 | 1.64 | Moderately soluble | No | No | No | No | No | 0 | 0 | 0 | 0 |
| ***S*O2** | 223.29 | 1.67 | Soluble | Yes | No | No | No | No | 0 | 0 | 0 | 0 |
| ***S*C2** | 297.80 | 2.82 | Moderately soluble | Yes | No | No | No | No | 0 | 0 | 0 | 0 |
| ***Se*O3** | 319.06 | 1.13 | Soluble | No | No | No | No | No | 0 | 0 | 0 | 0 |
| ***Se*C3** | 393.57 | 2.26 | Moderately soluble | No | No | No | No | No | 0 | 0 | 0 | 0 |
| ***S*O3** | 272.16 | 2.19 | Soluble | Yes | No | No | No | No | 0 | 0 | 0 | 0 |
| ***S*C3** | 346.67 | 3.44 | Moderately soluble | Yes | No | No | No | No | 0 | 0 | 0 | 0 |
| ***Se*O4** | 270.19 | 0.50 | Soluble | No | No | No | No | No | 0 | 0 | 0 | 0 |

| | | | | | | | | | | | | |
|---|---|---|---|---|---|---|---|---|---|---|---|---|
| ***Se*C4** | 344.70 | 1.64 | Moderately soluble | No | No | No | No | No | 0 | 0 | 0 | 0 |
| ***S*O4** | 223.29 | 1.67 | Soluble | Yes | No | No | No | No | 0 | 0 | 0 | 0 |
| ***S*C4** | 297.80 | 2.82 | Moderately soluble | Yes | No | No | No | No | 0 | 0 | 0 | 0 |
| ***Se*O5** | 285.16 | -0.12 | Soluble | No | No | No | No | No | 0 | 0 | 0 | 0 |
| ***Se*C5** | 359.67 | 1.01 | Moderately soluble | Yes | No | No | No | No | 0 | 0 | 0 | 0 |
| ***S*O5** | 238.27 | 0.91 | Soluble | Yes | No | No | No | No | 0 | 0 | 0 | 0 |
| ***S*C5** | 312.78 | 2.19 | Moderately soluble | Yes | Yes | No | No | No | 0 | 0 | 0 | 0 |
| ***Se*O6** | 276.14 | 1.16 | Soluble | No | No | No | No | No | 0 | 0 | 0 | 0 |
| ***Se*C6** | 350.65 | 2.29 | Moderately soluble | No | No | No | No | No | 0 | 0 | 0 | 0 |
| ***S*O6** | 229.25 | 2.29 | Soluble | Yes | No | No | No | No | 0 | 0 | 0 | 0 |
| ***S*C6** | 303.76 | 3.47 | Moderately soluble | No | No | No | No | No | 0 | 0 | 0 | 0 |
| ***Se*O7** | 300.22 | 0.46 | Soluble | No | No | No | No | No | 0 | 0 | 0 | 0 |
| ***Se*C7** | 374.72 | 1.59 | Moderately soluble | No | No | No | Yes | No | 0 | 0 | 0 | 0 |
| ***S*O7** | 253.32 | 1.70 | Soluble | Yes | No | No | No | No | 0 | 0 | 0 | 0 |
| ***S*C7** | 327.83 | 2.76 | Moderately soluble | Yes | Yes | No | No | No | 0 | 0 | 0 | 0 |
| ***Se*O8** | 246.19 | 0.43 | Very soluble | No | No | No | No | No | 0 | 0 | 0 | 0 |

| | | | | | | | | | | | | |
|---|---|---|---|---|---|---|---|---|---|---|---|---|
| ***Se*C8** | 320.70 | 1.68 | Soluble | No | No | No | No | No | 0 | 0 | 0 | 0 |
| ***S*O8** | 199.30 | 1.53 | Very soluble | Yes | No | No | No | No | 0 | 0 | 0 | 0 |
| ***S*C8** | 273.81 | 2.86 | Moderately soluble | No | No | No | No | No | 0 | 0 | 0 | 0 |
| ***Se*O9** | 296.25 | 1.41 | Soluble | No | No | No | No | No | 0 | 0 | 0 | 0 |
| ***Se*C9** | 370.76 | 2.61 | Moderately soluble | No | Yes | No | No | Yes | 0 | 0 | 0 | 0 |
| ***S*O9** | 249.36 | 2.63 | Soluble | Yes | Yes | Yes | No | No | 0 | 0 | 0 | 0 |
| ***S*C9** | 323.86 | 3.79 | Moderately soluble | Yes | Yes | Yes | No | No | 0 | 0 | 0 | 0 |
| ***Se*O10** | 280.18 | 0.85 | Soluble | No | No | No | No | No | 0 | 0 | 0 | 0 |
| ***Se*C10** | 370.74 | 2.35 | Moderately soluble | Yes | No | No | Yes | Yes | 0 | 0 | 0 | 0 |
| ***S*O10** | 233.29 | 2.03 | Soluble | Yes | Yes | No | No | No | 0 | 0 | 0 | 0 |
| ***S*C10** | 307.80 | 3.22 | Moderately soluble | Yes | Yes | No | No | No | 0 | 0 | 0 | 0 |
| ***Se*O11** | 298.29 | 1.32 | Soluble | No | No | No | No | No | 0 | 0 | 0 | 0 |
| ***Se*C11** | 336.33 | 2.13 | Soluble | No | No | No | Yes | No | 0 | 0 | 0 | 0 |
| ***S*O11** | 239.38 | 2.62 | Soluble | No | No | Yes | No | No | 0 | 0 | 0 | 0 |
| ***S*C11** | 325.90 | 3.68 | Moderately soluble | No | Yes | Yes | No | No | 0 | 0 | 0 | 0 |
| **Pentamidine** | 340.42 | 2.72 | Soluble | No | No | Yes | Yes | No | 0 | 0 | 0 | 0 |
| **Bortezomib** | 453.33 | 1.26 | Soluble | No | Yes | No | Yes | Yes | 0 | 0 | 1 | 0 |

[a]MW: Molecular weight

[b]Log P: The partition coefficient between n-octanol and water (log $P_{o/w}$) is the classical descriptor for lipophilicity. It is expressed as the arithmetic mean of the values predicted by the five proposed methods, described in Daina, A., *et al.* (Sci Rep 7, 42717 (2017)).

[c]Solubility: calculated according to ESOL model (J. Chem. Inf. Model. 44, 1000–1005 (2004).)
